# Membrane-binding properties of NS1 proteins from Zika and Dengue viruses: Comparative simulations in explicit bilayers reveal significant differences

**DOI:** 10.1101/2023.01.12.523767

**Authors:** Rajagopalan Muthukumaran, Ramasubbu Sankararamakrishnan

**Affiliations:** Department of Biological Sciences and Bioengineering, Indian Institute of Technology Kanpur, Kanpur 208016, India; Mehta Family Centre for Engineering in Medicine, Indian Institute of Technology Kanpur, Kanpur 208016, India

## Abstract

NS1 in flaviviruses is the only non-structural protein that is secretory and interacts with different cellular components. NS1 is localized in endoplasmic reticulum as a dimer to facilitate the viral replication. The crystal structures of NS1 homologs from zika (ZIKV) and dengue (DENV) viruses have revealed the organization of different domains in NS1 dimers. The β-roll and the connector and intertwined loop regions of wing domains of NS1 have been shown to interact with the membranes. The membrane-binding properties and the differences between ZIKV and DENV NS1 homologs in interacting with the membranes have not been investigated. In this study, we have performed molecular dynamics (MD) simulations of ZIKV and DENV NS1 systems in apo and in POPE bilayers with different cholesterol concentrations (0, 20 and 40%). In the simulations with bilayers, the NS1 protein was placed just above the membrane surface. At the end of 600 ns production runs, ZIKV NS1 inserts deeper inside the membrane compared to the DENV counterpart. The conformational landscape sampled by NS1 in the presence of membrane was analyzed. Unlike ZIKV NS1, the orientation of DENV NS1 is asymmetric in which one of the chains in dimer interacts with the membrane while the other is exposed to the solvent. The β-roll region in ZIKV NS1 penetrates beyond the headgroup region and some residues interact with the lipid acyl chains while the C-terminal region barely interacts with the headgroup. Specific residues in the intertwined region deeply penetrate inside the membrane with less interactions with water molecules. Our analysis showed that more charged residues of ZIKV NS1 are involved in stronger interactions with the headgroups than that found for DENV NS1. The role of hydrophobic and aromatic residues in interactions with acyl chain region is also evident. Presence of cholesterol affects the extent of insertion in the membrane and interaction of individual residues. This study clearly shows that the binding, insertion and interaction of ZIKV NS1 with the lipid bilayer significantly differs from its counterpart in DENV.

## Introduction

The flavivirus belongs to the family of *flavivridae* comprising ∼70 important human pathogens expressing high morbidity and mortality rates. Among these, the most important viruses that cause severe diseases in humans are dengue virus (DENV), West-Nile virus (WNV), Japanese Encephalitis virus (JEV) and recently, the Zika virus (ZIKV) ^1-4^. Most of these flaviviral infections are vector borne, primarily mosquitoes acting as carriers to human host. The major risk with ZIKV is the mode of transmission since the virus has adopted to non-insect vector routes like sexual transmission, blood and platelet transfusion as well as various body fluids, making it more vulnerable ^5, 6^. The primary symptom of DENV infections is a flu-like illness, which, in severe cases might lead to dengue haemorrhagic fever (DHF) or dengue shock syndrome (DSS) ^7^. In the case of ZIKV infection, it presents as a rash, flu-like illness and rarely results in Guillain–Barre syndrome in adults, with increasing evidence for neurological abnormalities in developing foetuses ^8^. ZIKV has gained global attention in the recent years. Since its major outbreak in 2007 and in 2016, the World Health Organization declared ZIKV as “a public health emergency of international concern”. It is alarming to note that ZIKV infections have been reported in ∼66 countries, including India ^9-12^.

Flavivirus is an enveloped virus with (+) ss RNA and a genome size of ∼11-12 kb. The genome is made up of a single open reading frame (ORF) flanked by 5’ and 3’ untranslated regions ^13^. This ORF is translated into a polyprotein, broken down to 10 structural and non-structural proteins. The three structural proteins (C, prM and E) form the envelope and basic skeleton of the virus, while the seven non-structural proteins (NS1, NS2A, NS2B, NS3, NS4A, NS4B and NS5) are responsible for replication and pathogenicity ^4, 14^. Interestingly, among all the proteins expressed by the virus, NS1 is the only secretory protein (sNS1) and hence acts as a biomarker to identify flaviviral infection. NS1 interacts with different host cell components and aids viral growth by evading normal host cellular processes ^15^. NS1 can occur in three different oligomeric states based on its function. In the dimeric state, it is localized in the ER lumen to aid in viral replication and three dimers associate with lipids to form a hexameric lipoprotein to interact with the cell surface receptors ^4^.

The mature viral NS1 is ∼350 amino acids long and exists in dimeric as well as hexameric states ^16, 17^. The crystal structures of DENV ^18^, WNV ^18^, and ZIKV ^19, 20^ have been solved and they all possess an overall similar structural fold. The monomeric NS1 contains three domains, namely, an N-terminal β-hairpin domain (residues 1-30), an epitope-rich wing domain (residues 31–181) and a C-terminal β-ladder domain (residues 182–352) with two potential glycosylation sites at N130 and N207. There are twelve invariant cysteine residues forming six disulfide bonds per monomer and are responsible for intra-domain stabilization. The β-hairpin domain forms the dimer interface between two monomers and intertwines to form a roll-like structure called as “β-roll” dimerization domain. The wing domain can be further divided into three subdomains, namely (i) α/β subdomain (residues 38–151), (ii) a long-intertwined loop (residues 91-130) and (iii) a discontinuous connector domain (residues 30-37 and 152-180) which connects the wing domain with the β-roll and C-terminal β-ladder domain. Residue numbering followed here is according to the Zika virus NS1 protein structure with PDB ID: 5K6K.

The β-ladder domain is made up of ten β-strands and upon dimerization, the 20 β-strands (10 from each monomer) are arranged like the rungs of a ladder (protein-protein interaction side) ^21^. The opposite surface of the ladder lacks a definite secondary structure and formed of loops, including spaghetti loops. Upon dimerization, NS1 has two faces, the inner face formed by the β-roll, second half of the intertwined loop, and β-ladder domain and the outer face formed by the spaghetti loop and first half of the wing domain ^16, 19, 21^. In the inner face, the β-roll, the connector subdomain and second half of the intertwined loop (108-130 a.a) forms a hydrophobic surface which will be favourable for the membrane interaction ^19, 20^. But the two flexible loops of wing domain, the intertwined loop (residues 108-130) and the finger loop (residues 159-163) regions are not resolved in the DENV (PDB ID: 4O6B) and WNV (PDB ID: 4OIE) crystal structures ^18^. However, in the recently solved ZIKV NS1 structure (PDB ID: 5K6K), these regions are visible as unstructured loops ^19^. As the intertwined loop is flexible, it is speculated to be involved in the interactions either with membrane or other proteins ^20^. It is reported that after translation, a part of NS1 is localized to the ER membrane (lumen side) to help in viral replication ^18-22^. Recently, an electron microscopy study on NS1 (ZIKV and DENV) explored its membrane binding property and has shown the role of NS1 in the formation of tubules protruding from the liposomal surface and inducing negative curvature which possibly hints the remodelling of ER membrane (formation of replication compartment)^23^. The mutational studies have shown the loss of liposome binding activity upon the substitution of residues from the finger loop (159-163), and the intertwined loop (W115, W118 and F123) in the NS1 protein ^19, 21^. It is also reported that NS1 interacts with the lipid rafts at the plasma membrane (which are highly enriched with cholesterol and sphingolipids). These lipid rafts are reported as preferred interaction sites for several viruses and its components ^22^.

NS1 is present in all flaviviruses and performs similar function across the entire family. However, NS1 homologs are sequentially diverse. The two major human pathogens DENV and ZIKV share a sequence identity of ∼50%, which affects the surface charge distribution, while having a similar three-dimensional structure (RMSD of 0.74 Å) ^20, 24^. A comparative study of DENV, WNV and ZIKV displays differences in electrostatic profiles which could have impact on the interactions of NS1 with partners like membrane, antibody, cellular receptors ^20, 25^. Apart from being a potential marker for disease identification, NS1 is considered as one of the important targets in anti-viral therapy. Several mutagenesis studies have explored the structure and function of NS1 domains ^26^.

A few structural and computational investigations on dimeric NS1 have recently been published. In the DENV NS1, a single amino acid substitution from proline to leucine at position 250 resulted in the loss of dimerization and a reduction in viral proliferation and pathogenicity. As a result, Oliveira et al used molecular dynamics simulations to study how a single residue change affects dimer stability. Their research reveals conformational alterations and a local network of interactions driven by a single interface residue mutation ^27^. Similarly, Roy et al. investigated the importance of disulphide bonds in maintaining dimeric unit stability and β-ladder domain structure using extensive molecular dynamics simulations and thermodynamic analyses ^28^. In addition to the structural stability of dimeric NS1, small molecule and peptide inhibitors targeting NS1 have been reported. A library of therapeutic compounds targeting ZIKV NS1 dimerization site was investigated by Raza et al. using molecular docking and molecular dynamics ^29^. Their study revealed top three potential compounds/molecules binding/interacting to the dimerization site at the β-roll motif residues ^29^. Songprakhon et al., on the other hand, employed ELISA to screen DENV NS1 against a peptide library, followed by molecular docking and molecular dynamics simulations ^30^. Although structural and inhibitory research on ZIKV and DENV have been published, the membrane binding property of NS1 is yet to be explored. Therefore, the current work employs all-atom molecular dynamics simulations to demonstrate the membrane binding property of NS1 and its possible association with cholesterol. For this purpose, we used homogenous POPE (1-Palmitoyl-2-oleoyl-sn-glycero-3-phosphatidylethanolamine) bilayer and two different cholesterol concentrations (20% and 40%) since it is one of the major components of lipid raft. Our results show that ZIKV NS1 inserts deeper inside the membrane, interacts with different components of the membrane and has stronger interactions with the cholesterol containing bilayers than that found for DENV NS1. These studies will help to have a deeper insight into the mechanism of viral infection and the regions of NS1 proteins that can be targeted by small molecules.

## Methodology

### NS1 protein apo system preparation and simulation

The NS1 crystal structures with PDB ID 5K6K (ZIKV) and 4O6B (DENV) were used as starting structures for MD simulations. The intertwined loop sections (residues 108-128) and finger loop residues (residues 159-165) are missing from the DENV crystal structure. As a result, the ZIKV crystal structure was used as a template to model these regions (ZIKV and DENV NS1 sequence share sequence identity of ∼50%). The GROMACS 5.1 software package was used to perform Molecular Dynamics (MD) simulations ^31^. The structures solvated with TIP3P water model ^32^ in a periodic box extended up to 12 Å from edge of the protein on all sides. The systems were neutralized by 10 and 6 Na+ ions respectively for ZIKV and DENV NS1 proteins. The systems were minimized using the steepest descent and conjugate gradient approaches. The minimized structures served as initial structures for the MD simulations. The temperature of the systems was maintained at 310 K by V-rescale coupling ^33^. The isothermal-isobaric ensemble was used to set the pressure to 1 bar and maintained by Parrinello-Rehman barostat ^34^. The short-range electrostatic interactions were treated with a cut-off value of 12 Å. The long-range electrostatic interactions were treated with the particle-mesh Ewald method. The systems were then equilibrated for 500 ps with harmonic restrictions on protein atoms utilizing NVT and NPT ensembles. After the equilibration, the production runs of the systems in which no restraints were applied were run for a period of 600 ns each.

### Preparation of NS1-model membrane Systems

The NS1 protein was manually placed over the pre-equilibrated lipid bilayer just above the lipid head group (distance between protein centre of mass and lipid head group is ∼21 Å, three-dimensional structure is shown in Figure 1). Details of the preparation of pre-equilibrated lipid bilayer are provided in Supplementary Information. The overlapping lipid molecules are removed to avoid the steric clashes (∼12-15 lipid molecules removed). The protein along with the bilayer was minimized using steepest descent and conjugate gradient methods. The system was solvated with TIP3P ^32^ water molecules in a periodic box with dimensions of 130×130×150 Å^3^, ensuring at least 12 Å of water molecules between the protein and box edge. The system was neutralized with sodium ions by replacing solvent molecules (Figure 1) and was then equilibrated with harmonic positional restraints on the protein (backbone and side chain atoms) and lipid bilayer headgroup atoms. The restraints were gradually removed in steps of 500 ps. Berendsen temperature and pressure coupling methods were applied during this process ^35^. The temperature of the system was maintained at 310 K by V-rescale coupling ^33^. The pressure was set to 1 bar using the semi-isotropic coupling scheme to a parrienello-Rahman pressure coupling baraostat ^34^. The short-range interactions were truncated with a cut-off of 12 Å. The Long-range electrostatic interactions were treated with the particle-mesh Ewald method ^36^. The system was equilibrated for 50 ns without restraints followed by a production run of 600 ns using the same simulation protocol described above. GROMACS modules like (rms, rmsf, density and do_dssp) were used to analyze Root Mean Square Deviation (RMSD), Root Mean Square Fluctuation (RMSF), density plots and secondary structures ^31^. The electrostatic potentials were generated by solving Poisson-Boltzmann equation utilizing Adaptive Poisson-Boltzmann Solver (APBS) ^37, 38^ server and visualized using Chimera^39^. The interaction energies between NS1 and membrane were calculated by summing up the van der Waals and electrostatic energy terms with a cut-off distance of 12 Å. The complexes are denoted as ZIKV_APO_, DENV_APO_ for NS1 without membrane, ZIKV_POPE_ and DENV_POPE_ for NS1 in complex with POPE neat bilayers, and ZIKV_CHOL20_, ZIKV_CHOL40_, DENV_CHOL20_ and DENV_CHOL40_ for the systems in which POPE with 20% and 40 % of cholesterol concentrations were used. A total of 8 systems were simulated and the simulation details are summarized in Table. 1.

**Figure 1:**
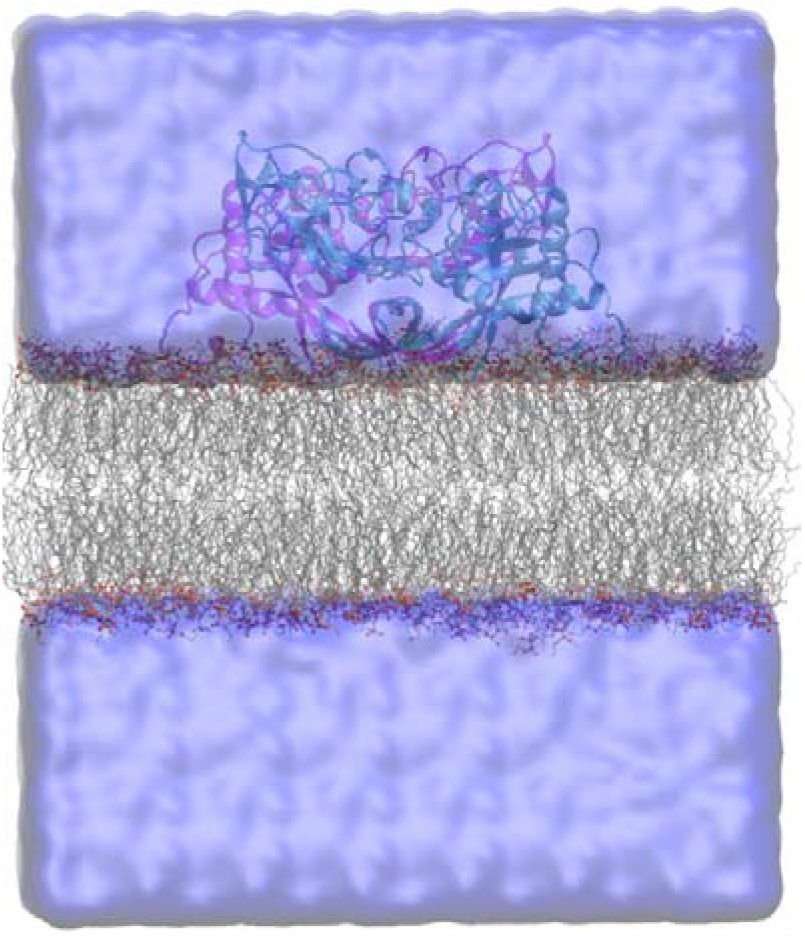
Initial simulation setup of NS1-membrane complex. The two chains of NS1 are shown as cartoon representations in cyan and magenta colors. The membrane is shown in stick representation (grey color) and water molecules as surface representation. The following color convention was used: carbon: turquoise, nitrogen: blue, oxygen: red and hydrogen: white.

**Table 1:** Summary of simulations.

| System | Type of bilayer | System | Simulation time (ns) |
| --- | --- | --- | --- |
| **ZIKV** | NS1 apo | ZIKV<sub>APO</sub> | 600 |
|  | POPE | ZIKV<sub>POPE</sub> | 600 |
|  | POPE with 20% cholesterol | ZIKV<sub>CHOL20</sub> | 600 |
|  | POPE with 40% cholesterol | ZIKV<sub>CHOL40</sub> | 600 |
| **DENV** | NS1 apo | DENV<sub>APO</sub> | 600 |
|  | POPE | DENV<sub>POPE</sub> | 600 |
|  | POPE with 20% cholesterol | DENV<sub>CHOL20</sub> | 600 |
|  | POPE with 40% cholesterol | DENV<sub>CHOL40</sub> | 600 |

## Results

### NS1 Sequence variation

The flaviviral NS1 is the only viral protein that is secretory in nature and engaged in several functions, including viral reproduction, immune evasion, and pathogenesis ^26^. The dimeric NS1 anchors to the ER lumen and assists in the development of the replication compartment ^21, 23^. The goal of the study was to characterize interactions of ZIKV and DENV NS1 protein with membrane and how they are affected by different membrane compositions as a function of cholesterol concentration. We manually placed ZIKV and DENV NS1 above (distance between protein centre of mass and lipid head group is ∼21 Å) the pre-equilibrated bilayers and 600 ns of all-atom molecular dynamics were performed to evaluate the membrane binding properties of NS1. The results are analyzed to find out how NS1 proteins interact with the membrane and any potential differences in the mode of membrane binding and interactions between the NS1 homologs of ZIKV and DENV.

Although ZIKV and DENV NS1 homologs show sequence variation, biochemical analysis reveals that they have similar properties ^18, 19^. The most important property of NS1 is membrane binding and membrane remodelling for viral replication compartment^23^. As the protein first encounters the lipid headgroups when binding to membranes, it is important to know the charge distribution between the two NS1 homologs. To explore this, we performed structure-based sequence alignment using ZIKV crystal structure as reference (PDB ID: 5K6K (Figure S2). The analysis of structure-based sequence alignment reveals frequencies of basic and aromatic residues are greater in the ZIKV NS1 homolog than in DENV NS1. The β-roll and wing domains (connector, intertwined loop regions) have been recognized to mediate contact with the lipid bilayers ^19, 20, 40^. Hence, we focused our analysis on these domains, particularly the inner face area. In the wild-type sequences, the residues N10, T29 of β-roll domain, Q31, Q35, Y158, T164 of connector domain, S40, A99, A141 of α/β subdomain, S103, Y113, S125, S128, of intertwined loop and N191, L206, E213, S216, T265, V286, E326, L338, N347 of C-terminal domain in DENV NS1 are substituted with basic amino acids in ZIKV NS1 (the residues numbering is with respect to ZIKV NS1 crystal structure 5K6K). Similarly, the residues N9 and E26 of the β-roll domain, Q48, S58, A77, and Q85 of the α/β subdomain, P101, Q102, P105, P112, Y122, and T129 of the intertwined loop, and A182, E192, E273, S323, and P341 of the C-terminal domain of ZIKV NS1 are replaced with basic amino acids in DENV NS1. In total, 22 residues of DENV and 17 residues of ZIKV NS1 proteins from β-roll and wing domain are substituted by basic residues in the corresponding positions of ZIKV NS1 and DENV NS1 respectively (Figure S2). Also, residues T22, Q98, K122, M123 and Q175 in DENV NS1 and E178 and T256 residues of ZIKV NS1 are substituted by aromatic residues respectively in ZIKV and DENV NS1 homologs. Altogether, the inner face of ZIKV NS1 is dominated by basic residues especially the connector domain which plays significant role in the interaction with the membrane and to a lesser extent by aromatic residues to facilitate interaction with the lipid bilayer. All-atom molecular dynamics simulations were performed to find out the role of basic and aromatic residues responsible for stabilizing the NS1 over a lipid bilayer at atomic level.

### Conformational dynamics of NS1 over different lipid bilayers

The solvated NS1 homologs of ZIKV and DENV were initially simulated without lipid bilayers for 600 ns (ZIKV_APO_, DENV_APO_). The interaction of NS1 with membrane is modeled using pre-equilibrated bilayers of POPE with varying concentrations of cholesterol (0 %, 20 %, and 40 %). Considering that the NS1-membrane contact is a surface interaction driven by the inner face of NS1, the protein was placed on the membrane surface and simulated for 600 ns to allow protein insertion. For NS1 investigated in neat POPE bilayers, the systems are designated as ZIKV_POPE_ and DENV_POPE_. The stability of NS1 was assessed by measuring the root mean square deviations (RMSD) of backbone atoms of chain A and chain B. The ZIKV_POPE_ stabilized after 100 ns with minimal structural fluctuation at an average value of 2.06±0.19Å, whereas the DENV_POPE_ stabilized after 400 ns with an average value of 3.45±0.23 Å, suggesting overall structural stability (Figure 2a and 2b) for both systems, although DENV NS1 exhibits higher deviation. In contrast, the cholesterol-containing systems have a higher RMSD than the NS1 systems simulated with neat POPE bilayers. The ZIKV_CHOL20_ RMSD stabilized with an average value of 3.72±0.34 Å after 300 ns, whereas DENV_CHOL20_ RMSD stabilized with an average value of 3.18±0.21 Å after 150 ns. The complexes ZIKV_CHOL40_ and DENV_CHOL40_ showed a more dynamic behaviour as the RMSD increased up to 4-5 Å during the initial 350 ns and shows a reduced stabilization at an average RMSD of 3.62±0.80 Å and 3.95±0.44 Å respectively (Figure 2a and 2b). In general, the presence of unstructured loops (intertwined loop and finger loop) in NS1 may lead to an increase in RMSD thus, the RMSD calculations were redone after excluding these loops. Although a similar pattern of RMSD is observed, the deletion of these unstructured loops has a negligible impact on the global RMSD for all simulated systems, affecting it by just 1 Å (Figure S3a and S3b).

**Figure 2:**
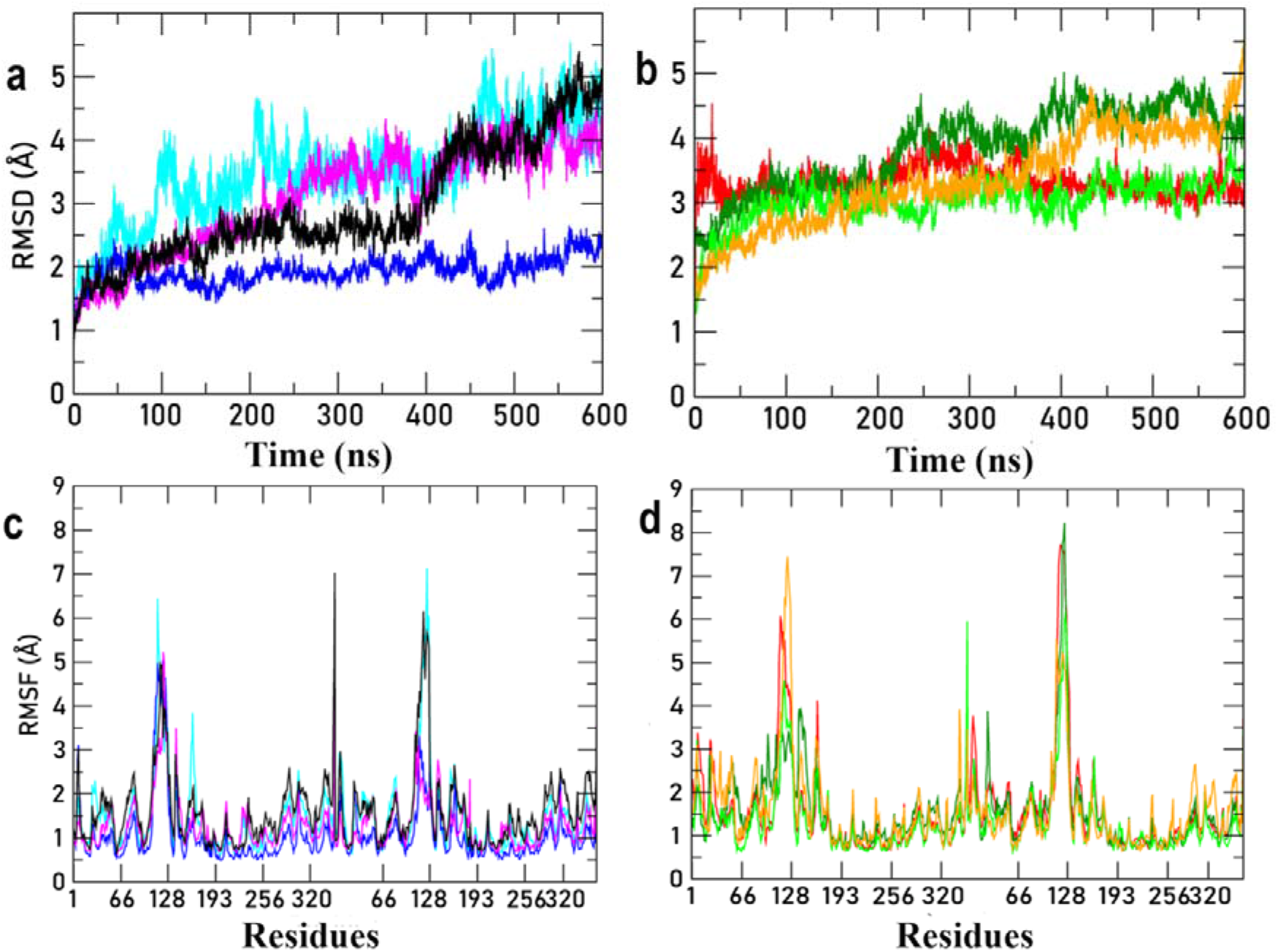
Time evolution of RMSD of ZIKV NS1 (a) and DENV NS1 (b) and RMSF of ZIKV NS1 (c) and DENV NS1 (d). RMSD was calculated for the backbone atoms of NS1 dimer (both Chains A and B) with respect to the starting structure. Only Cα atoms were considered to calculate RMSF values. The different NS1 complexes are differentiated as ZIKV_APO_ (cyan), ZIKV_POPE_ (Blue), ZIKV_CHOL20_ (magenta), ZIKV_CHOL40_ (Black), DENV_APO_ (Red), DENV_POPE_ (Dark green), DENV_CHOL20_ (light green) and DENV_CHOL40_ (orange).

In addition to the root-mean-square deviation, the root-mean-square fluctuation (RMSF) of the Cα atoms across the 600 ns simulation period was calculated and displayed in Figure 2c and 2d. Apart from few C-terminal residues (residues 350-352 of ZIKV_CHOL40_ and DENV_CHOL20_) and unstructured loop regions (residues 108-130), the RMSF exhibits a stable dynamics of the overall NS1 structure. The intertwined loop residues (residues 108-130) of dimeric NS1 have an additional flexibility of up to 5 Å. The intertwined loop displays an unequal pattern of flexibility, with one chain exhibiting greater fluctuation than the other (for instance, the chain A of ZIKV_POPE_ shows a fluctuation of ∼5 Å whereases chain B shows 3 Å). Notably, the hydrophobic finger loop residues (residues 155-170) exhibited higher variations in the absence of membrane (3.9 Å) than in membrane-bound complexes (3.0 Å forDENV and 2.0 Å for ZIKV complexes), indicating that the loop is stabilized by the membrane interactions (Figure 2c and 2d).

### NS1 orientation at the membrane

It is known that one of the key functions of NS1 is to interact with lipid bilayers and lipid rafts. The interaction is mediated by the inner face of NS1 comprised of the extended hydrophobic projections of the intertwined loop and finger loop of the wing domain ^19, 20, 23^. In our study, the interaction of NS1 with the lipid membrane was examined by manually placing the NS1 over the membrane in such a way that the distance between protein centre of mass and lipid head group is ∼21 Å. In general, a peripheral membrane protein can explore configurational dynamics by either inserting into the membrane or exploring several orientational arrangements on the surface of the membrane. The dynamic properties of NS1 over the membrane were analysed in our study by examining the distance and angle of inclination with respect to the membrane. The distance *(d)* was monitored between the center of mass of all heavy atoms of NS1 (COM_p_) and the membrane (COM_M_). The angle (tilt) is derived as the angle between the vectors linking the C-terminal residues 321-326 (β17; Figure S2) of both monomers to the membrane normal vector (Z-axis). These two parameters were used to quantify the insertion and to establish the correlation between insertion depth and tilt angle. We have plotted the MD trajectories of the distance *d* and the angle θ as a function of time (Figure 3a and 3b). This analysis reveals the distribution of conformations sampled throughout the simulation, allowing for the identification of the most preferred combination (of distance versus angle).

**Figure 3:**
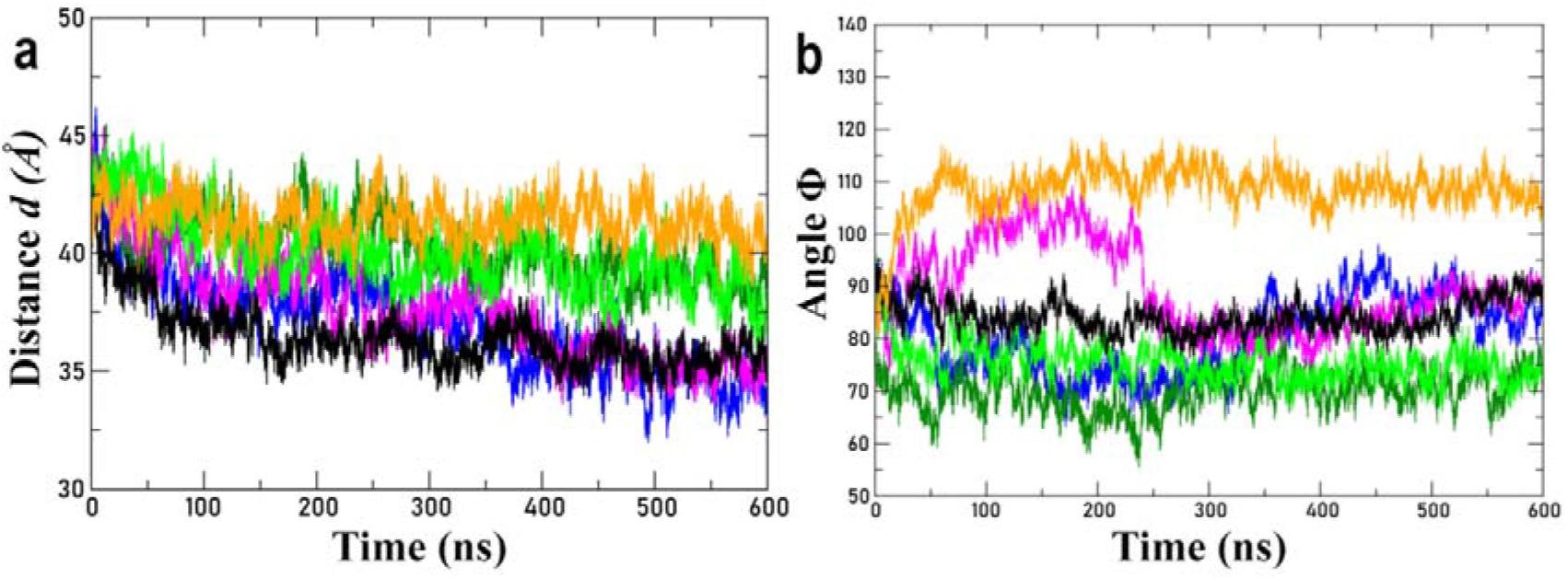
The orientation of NS1 with respect to the membrane plane plotted as a time evolution of distance *d* and angle θ. The different NS1 complexes are differentiated as ZIKV_POPE_ (Blue), ZIKV_CHOL20_ (magenta), ZIKV_CHOL40_ (Black), DENV_POPE_ (Dark green), DENV_CHOL20_ (light green) and DENV_CHOL40_ (orange).

We analyzed the time evolution of distance *d* and angle θ to find out if there is any relationship between insertion depth and binding orientation (Figure 3a and 3b). At the beginning of the production run, the distance *d* remains similar for all six systems (∼ 42.01± 1.1 Å) shown in Table 1. The reduction in *d* value implies that NS1 is inserted into the membrane, whereas the increase in distance will indicate that NS1 is moving away from the membrane’s center. As depicted in Figure 3a, a significant divergence in the dynamics of ZIKV and DENV NS1 has been observed. The time evolution of the distance *d* reveals that ZIKV NS1 inserts deeper inside the membrane than that observed for DENV NS1 and this difference is clear irrespective of cholesterol concentration. The insertion of ZIKV NS1 into all three membranes is quite comparable and there is significant overlap in the trajectories well after 350 ns with an average distance of 35Å (from the centre of the membrane), indicating a deeper insertion (as the starting distance is ∼42Å). However, DENV NS1 insertion is less deeper than ZIKV NS1. All three DENV NS1 maintained the same distance until 250 ns (42 Å), however after 250 ns, DENV_POPE_ and DENV_CHOL20_ NS1 demonstrate a reduction in value to 39 Å, which is at least 4 Å closer to the membrane-water interface from the center of the membrane than that found for the ZIKV NS1 (∼35 Å). In the case of DENV_CHOL40_, *d* remains close to the initial distance ∼42 Å (figure 3a). Although the initial distance *d* was same for all six simulations, the ZIKV complexes insert more deeper inside the membrane than that found for the DENV counterpart.

As the first contacts of NS1 with the membrane occur via the β-roll domain, finger loop and intertwined loop, it adopts various conformations during simulations and therefore, the tilt angle θ was determined using the C-terminal domain as a reference as mentioned earlier. MD trajectories of θ as a function of time are plotted for all six simulated systems (Figure 3b). At the beginning of the production run both ZIKV and DENV complexes (ZIKV_POPE_, DENV_POPE_, ZIKV_CHOL20_, DENV_CHOL20_, ZIKV_CHOL40_ and DENV_CHOL40_) remain at a θ value of 90°. During production run the time evolution of θ shows that the ZIKV systems tilt less than that found for DENV systems as ZIKV complexes attain a θ value of 90° towards the end of the 600ns simulation. Though the ZIKV NS1 stabilized at a θ value of 90°, the ZIKV_POPE_ and ZIKV_CHOL20_ showed a cyclic change in the tilt angle. In ZIKV_POPE_, the θ value decreases upto ∼70° (∼310 ns) followed by a gradual increase of θ value reaching 90° (Figure 3b). Similarly, in ZIKV_CHOL20_ the θ value increases upto ∼105° (for ∼240 ns) followed by a decrease in the tilt angle to 80° and gradually reaching 90°. In contrast to this observation, the DENV systems exhibit a different behavior and remain tilted at θ =75° (DENV_POPE_ and DENV_CHOL20_) and θ = 110° (DENV_CHOL40_) throughout 600 ns simulation time (Figure 3b). In summary, this analysis reveals that ZIKV NS1 protein inserts deeper inside the membrane with its overall orientation maintained parallel to the membrane surface. However, DENV NS1 remains relatively closer to the membrane-water interface and is tilted with respect to the plane of the membrane.

We have plotted *d* versus θ contour diagrams for the NS1 membrane complex systems to understand the conformational landscape sampled by the ZIKA (Figure S4 a-c) and DENV NS1 (Figure S4 d-f) systems in different cholesterol conditions for 600 ns. The contour map of ZIKV_POPE_ displays two orientation clusters, one of which is sampled at *d* = ∼35 Å and θ around 87-90° and the other at *d* = ∼38 Å and θ = 70-75° tilt (Figure S4a and S5a-c). A representative structure was extracted from each cluster of the ZIKV, DENV complexes and is depicted in Figure S5 and S6. The three-dimensional structure of ZIKV_POPE_ complex reveals that when the θ value is < 90°, the insertion of NS1 into the membrane is lower when compared to a θ value approaching 90° (Figure S5b-c). In the case of ZIKV_CHOL20_, three distinct clusters with one sparsely populated is observed, indicating that NS1 covering larger conformational space. The two well populated clusters are located at (*d* = 34-35 Å, θ = 84-90º) and (*d* = 37-38 Å, θ = 79-83°) and the sparsely populated cluster at (*d =* 38-40 Å, θ = 95-105°) (Figure S4b and figure S5d-g). It is observed that as the θ value is closer to 90°, a maximum insertion of NS1 into the membrane is established (Figure S5f). Interestingly, the ZIKV_CHOL40_ displays just one populated cluster with *d* = 34-38 Å and θ = 80°-90° (Figure S4c, S5h-i).

In contrast to the ZIKV systems, the contour maps for all three DENV systems only display a single populated cluster (Figure S4d-f). The sample range for DENV_POPE_ is confined to *d* = 38-42 Å and θ = 60-75° (Figure S4d). Presence of cholesterol favors the insertion of DENV NS1 marginally, as seen by the contour map. Similar to the DENV_POPE_ system, DENV_CHOL20_ and DENV_CHOL40_ sample at *d* = 38-42 Å (Figure S4 e,f). However, the C-terminal domains of NS1 in both these systems are tilted in opposing directions with respect to membrane plane. In DENV_CHOL20_, the θ value fluctuates between 70° and 80°, whereas in DENV_CHOL40_, it varies between 105° and 110°(Figure S4e,f). Upon visualizing the representative structures, the binding mode of DENV NS1 is analogous to that of ZIKV NS1 with variation in the insertion and angles (Figure S6a-f). The ZIKV and DENV complex establish a clear link between the two parameters *d* and θ, where deeper insertion results in a balanced NS1 that permits the C-terminal domain to interact with the membrane.

### Binding of NS1 domains to membranes

The analysis of *d* and θ confirms that both ZIKV and DENV NS1 interacts with the membrane during simulation, although the degree of insertion varies. To quantify the insertion of NS1 and its distinct domains, the density distribution of different domains of NS1, lipid headgroup, glycerol and cholesterol atoms (PO4 for POPE and O for cholesterol) of the interacting layer (one of the two layers) along the z-axis (perpendicular to the bilayer) was computed. The density distribution is plotted in Figure 4 and S7 where the zero in x-axis corresponds to the center of the lipid bilayer. In these figures, the grey filled band depicts the density of PO4 head group and the thick black line identifies the peak. The maximal density of the lipid headgroup (PO4) of POPE, 20% and 40% cholesterol containing bilayers reside at 21.5 Å, 22 Å, and 22.9 Å, respectively. Similarly, the highest density for cholesterol O3 atoms in these systems occurs at 19.5 Å (the cyan coloured area in the Figure 4 and S7). When the density peak of the NS1 domain overlaps with the density peak of the lipid PO4 group or if it goes towards the center of the membrane, we can interpret that the domain is buried beneath the corresponding lipid group and reflects the distance from the lipid bilayer center.

**Figure 4:**
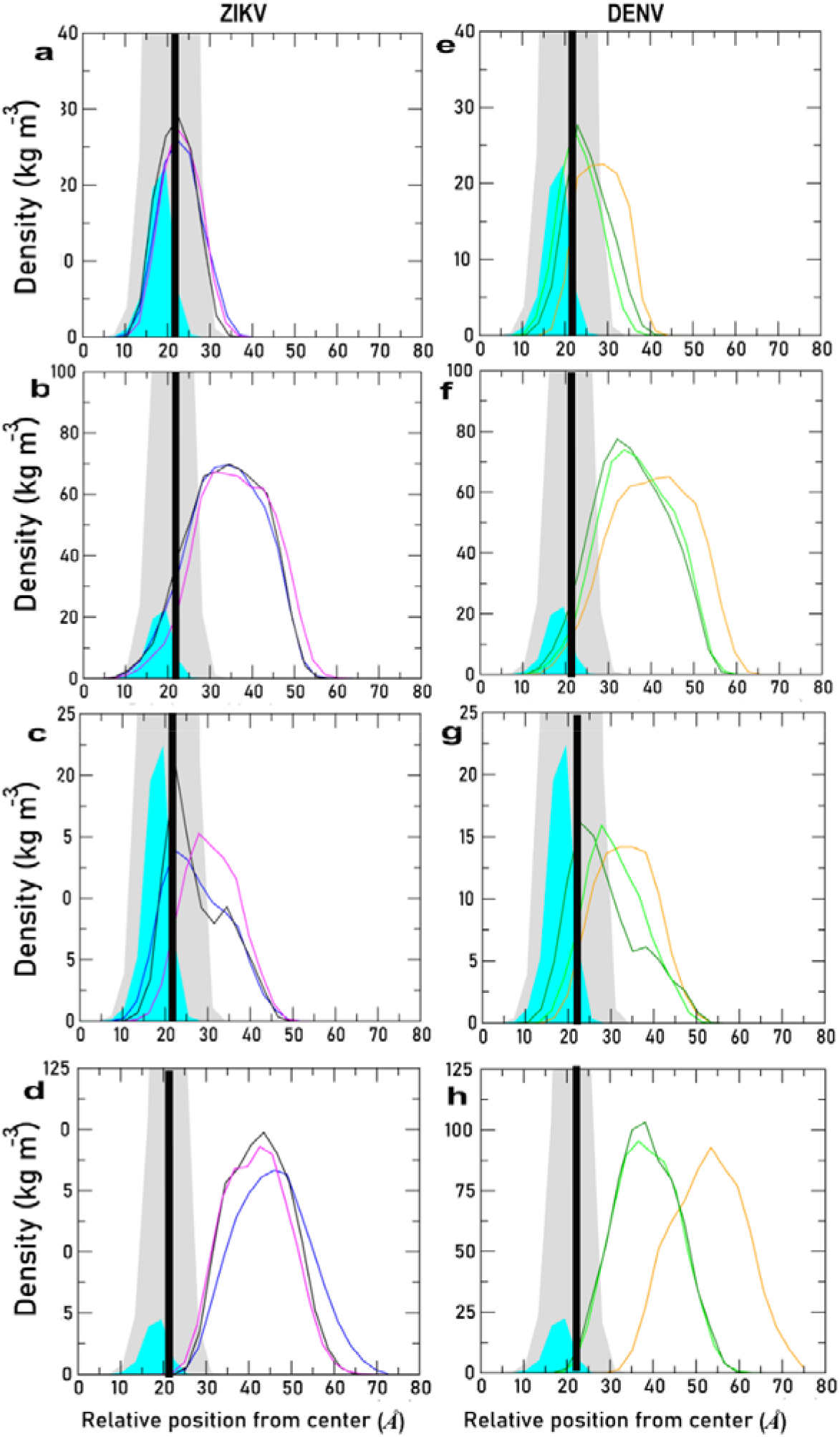
Average density profiles calculated for the individual domains of NS1 (Chain A) simulated over the three different bilayers along the Z-axis of the bilayer. The different NS1 complexes are differentiated as ZIKV_POPE_ (Blue), ZIKV_CHOL20_ (magenta), ZIKV_CHOL40_ (Black), DENV_POPE_ (Dark green), DENV_CHOL20_ (light green), and DENV_CHOL40_ (orange). The lipid headgroup region is shown in the grey background with the peak position indicated by the thick black line. The cholesterol density shown in the cyan. The zero on the x-axis corresponds to the bilayer center (as NS1 interacts with top layer and hence only top layer is shown in the figure). The profiles for individual domains are shown in rows labelled β-roll (a,e), wing (b,f), Intertwined loop (c,g) and the C-terminal domain (d,h) for chain A of ZIKV, DENV NS1 proteins.

The density profiles calculated for individual domains of both chains of dimeric NS1 across the three distinct bilayers (ZIKV_POPE_, ZIKV_CHOL20_, ZIKV_CHOL40_ in blue, magenta and black colors respectively) are shown in the Figure 4. Understanding the NS1 interaction pattern is enhanced by the density distribution of individual domains. The peak density of the β-roll domain (22.2 Å) of both chains overlap well with the peak density of the phosphate headgroup, glycerol group and enables to interact well with the acyl groups (Figure 4a and S7a). Majority of the wing domain remains exposed to the solvent and only a small portion of the wing domain, including the finger and intertwined loop, interacts with the membrane. The wing domain’s peak density (at 34 Å) remains 12 Å away from the lipid headgroup towards the aqueous medium and exhibits less overlap with the headgroup density profile (Figure 4b and S7b). By spanning its peak density over the headgroup, glycerol, the uncharacterized intertwined loops (of the wing domain) are well embedded into the membrane (Figure 4c and S7c). This loop also exhibits a little insertion asymmetry between the two monomers, with chain B inserted deeper (Z=19.0 Å) than the chain A (Z=22.2 Å). Major part of the C-terminal domain remains away from the lipid headgroup region (Figure 4d and S7d).

The NS1 density distribution of the ZIKV_CHOL20_ complex (Figure 4, magenta) is comparable to that of the ZIKV_POPE_ complex (shown in blue). The peak density of β-roll domain is situated at Z=22.0 Å, overlapping the phosphate headgroup and partly with cholesterol O3 atoms (Figure 4a and S7a, magenta). An asymmetric insertion of intertwined loops was observed in ZIKV_POPE_, which is more prominent in ZIKV_CHOL20_ as the peak densities stay around Z=27.9 Å for Chain A and Z=22.0 Å for Chain B. As a result, chain B intertwined loop possibly interact with more cholesterol atoms than chain A due to more overlap with cholesterol head group (Figure 4c and S7c, magenta). The C-terminal density profile is similar to that of the ZIKV_POPE_ complex (Figure 4d and S7d, blue and magenta). The ZIKV_CHOL40_ system exhibits marginally more insertion than the ZIKV_POPE_ and ZIKV_CHOL20_ systems, as seen by overlapping lipid headgroups. The maximal density of the β-roll domain occurs at Z= 22.5 Å (the head group phosphate is located at 22.9 Å), which is well within the interaction range of acyl chains (Figure 4a and S7a, black). In comparison to ZIKV_POPE_ and ZIKV_CHOL20_ complexes, the intertwined loop shows a nearly symmetrical insertion profile (Z=22.5 Å for both chains) and overlapping with cholesterol density (Figure 4c and S7c, black).

Similar to ZIKV complexes, density profiles were plotted for individual domains of DENV complexes (Figure 4 and S7). The density profile for β-roll domain is asymmetric (peak: 22.9 Å, 25.9 Å) despite its overlap with the phosphate headgroup (density lies at 21.3 Å). Still the magnitude of insertion is less than that of ZIKV (∼4 Å difference less deeply inserted). The wing domain partially overlaps with the headgroup and glycerol sections, and much of the area is solvent exposed. The intertwined loop demonstrates an uneven distribution with a density peak at Z= 19.8 Å and 25.9 Å indicating that it is more towards the aqueous medium than that found for ZIKV_POPE_. Similar to ZIKV_POPE_, the C-terminal domain has an irregular insertion pattern for two chains, and it is completely exposed to the solvent (Figure 4d and S7d).

In the case of DENV_CHOL20_ (light green), the distribution follows a similar pattern to that of DENV_POPE_ (dark green) shown in the Figure 4e. The maximal densities of the β-roll domain are found at Z=21.9 Å for chain A and 24.8 Å for chain B, which overlaps with the phosphate headgroup, glycerol (Figure 4a and S7a). The β-roll domain of DENV_CHOL20_ inserts deeper inside the membrane than DENV_POPE_ but less than ZIKV_CHOL20_ (Figure 4a and S7a). Surprisingly, the intertwined loop has a symmetrical insertion distribution with a maximum density at Z= 27.8 Å (Figure 4c and S7c), which is close to ZIKV_CHOL20_ (chain A, Figure 4c).

In comparison to DENV_POPE_ and DENV_CHOL20_, the DENV_CHOL40_ has different density profiles as it inserts less into membrane (Figure 4 and S7, shown in orange). The maximum density of the β-roll domain lies at Z=27.4 Å for chain A, 22.9 Å for chain B (Figure 4a and S7a). The β-roll domain slightly overlaps with head group and primarily interacts with the phosphate headgroup, but also occasionally with the acyl group. The intertwined loop exhibits an asymmetrical density profile with peak lies at Z= 32.0 Å, 25.9 Å (Figure 4c and S7c), hence chain B interacts more effectively with the membrane headgroup region than the chain A, but to a lesser extent than DENV_CHOL20_ and ZIKV_CHOL40_ (Figure 4g and 4c). A major part of the C-terminal domain remains away from the lipid headgroup region with a highly asymmetrical profile (Figure 4d and S7d).

Analysis of density profiles reveals the degree of insertion of domains into distinct lipid components in membranes with different cholesterol concentrations. The β-roll domain of ZIKV complexes exhibits insertion inside the membrane (∼3.0 Å) relatively deeper than the corresponding DENV complexes. In most cases, the densities of β-roll, intertwined loop and finger loop overlap with the headgroup density and partly with cholesterol enabling the NS1 to interact with the hydrophobic regions of the lipid bilayer. The density plots and RMSF analyses (distinct fluctuations by the individual chains) reveal an asymmetric insertion profile for the intertwined loop, with one chain inserting more than the other. To understand this phenomenon further, we have analyzed the insertion nature of individual residues in detail.

### Conformational orientation of spike region at the membrane

The Intertwined loop has been implicated in membrane binding in both ZIKV and DENV ^20^, and this is the first study to evaluate interaction with a lipid membrane of variable compositions. The intertwined loop residues lack a definite secondary structure and is disordered. Due to its high flexibility, the loop structure was not observed in previously deposited crystal structures ^18, 21^. But the loop coordinates are revealed by the latest ZIKV NS1 crystal structure, although it lacks well-defined secondary structure ^19^. In this work, the intertwined loop of the ZIKV NS1 crystal structure was employed as a template to model the DENV NS1 loop. During simulations, the intertwined loop region (residues 108-130) effectively inserts into the bilayer as seen through density analysis and also the distribution of loop density demonstrates that a part of the loop overlaps with the density of lipid membrane components, indicating that this loop region interacts with the lipid headgroup (Figure 4c, S7c). As it interacts with the membrane, it may undergo structural changes throughout simulations and adopt different conformations. Consequently, the superimposition of snapshots extracted from the last 300 ns of simulation (at an interval of 20 ns) over the initial structure reveals two distinct conformations: (i) a stable membrane-inserted state and (ii) a solvent-exposed state with structural heterogeneity (Figure 5a-h).

**Figure 5:**
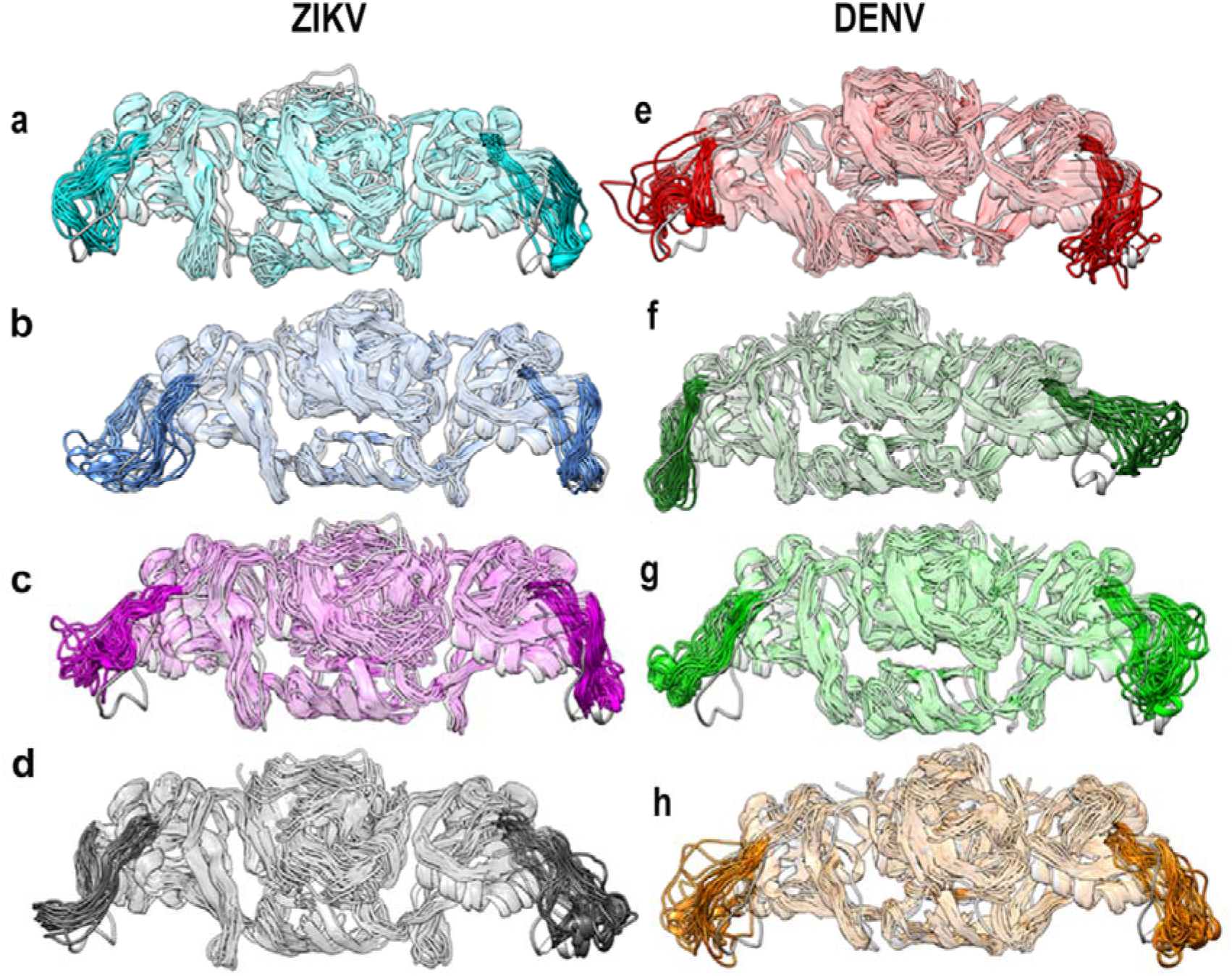
MD simulated NS1 structures, extracted over the last 300 ns at an interval of 20 ns, superposed on the initial structure (grey). The intertwined loop regions are highlighted, and other regions are shown as transparent. The different simulation systems are colored as, ZIKV_APO_ (cyan), ZIKV_POPE_ (blue), ZIKV_CHOL20_ (magenta), ZIKV_CHOL40_ (black), DENV_APO_ (red), DENV_POPE_ (dark green), DENV_CHOL20_ (light green) and DENV_CHOL40_ (orange). Structures are shown in ribbon representation.

The MD trajectory was investigated for the presence secondary structural changes and so the time evolution of the secondary structure for each amino acid in the intertwined loop was plotted using DSSP (shown in Figure S10 for chain A a-d and chain B e-h). In the apo form, residues 117-125 form β-bend, whereas the remaining regions of the intertwined loop are unstructured coil. In the case of ZIKV_POPE_, the residues 121-126 form a bend or 3_10_-helix (at various times of simulation) in chain A (Figure S10 Ib), while the same residues in chain B forms a turn structure (Figure S10 If). In cholesterol-containing systems, the intertwined loop often forms a β-turn or β-bend conformations as seen in the chain A of ZIKV_CHOL20,_ chain B of ZIKV_CHOL40_ (residues 113-118 and 119-125), chain B of ZIKV_CHOL20_ (residues 125-128) and ZIKV_CHOL40_ (118-127) (Figure S10c-d, h). Interestingly, the residues 120-126 of chain B in ZIKV_CHOL20_ show an α-helix conformation, a unique conformation among the ZIKV simulation systems (Figure S10g).

In DENV systems, however, such α-helical structures were more frequently observed. The chain A residues 118-125 DENV_APO_ remain in a coil, whereas the other chain displays a beta-turn conformation (Figure S10IIa,e). In DENV_CHOL20_, the chain A residues (119-125 a.a) forms β-turn while the same residues of chain B intermittently forms an α-helix till 450ns (Figure S10IIg). Similarly, the residues 118-125 in both chains adopt an α-helix of DENV_CHOL40_ complex (Figure S10h). Altogether, it is noted that a β-bend or β-turn structure is seen in the intertwined loops inserting into the membrane, while the solvent exposed loop is present as a highly flexible unstructured coil.

Our analysis indicates that the intertwined loop inserts into the membrane. Hence, we measured the average insertion depth of individual residues (distance between Cα atoms of each residue to COM_M_) across all simulated systems (Figure S11 and S12). As reference, the average position of the phosphate headgroup from COM_M_ is indicated by a red line and it is ∼21-22 Å from the COM_M_. The residues with average insertion depth values less than 21-22 Å imply that those residues are inserted deeper inside the membrane and is likely to be buried beneath the phosphate headgroup. During simulations, residues of the intertwined loop are buried to varying extents.

According to the density distribution discussed earlier, an asymmetric insertion has been noted in ZIKV_POPE_ and ZIKV_CHOL20_ systems while a symmetric insertion is seen in ZIKV_CHOL40_ (Figure 4 and S7, blue, magenta, and black). Consequently, the chain A residues 115-120 and chain B residues 115-123 of ZIKV_POPE_ and chain B residues 118-125 of ZIKV_CHOL20_ are inserted well into the membrane as seen in Figure S11a and S12a. The chain A residues of ZIKV_CHOL20_ remains above the level of headgroup (Figure S11a). In the case of ZIKV_CHOL40_, the density distribution of both chains demonstrates a symmetric insertion, and the residues 117-125 stayed close to the phosphate headgroup level (Figure S11a and S12a). The four intertwined loop conformations (ZIKV_CHOL40_, ZIKV_POPE_ and ZIKV_CHOL20_) which show insertion into the membrane have a common secondary structure, a β-bend or a β-turn (Figure S10).

The interaction with the membrane also affect their hydration state. Hence, the number of waters within 4 Å of each of the loop residues was calculated and plotted in Figure S11c and S12c. The polar and charged (especially basic) residues, such as K116, K120, S121, and R125, have a greater hydration state than hydrophobic residues, such as A117, G119, and V124. The general assumption of hydration state being directly proportional to insertion depth correlates only in case of aromatic residues like W115 and W118, which are above the headgroup and well hydrated in ZIKV_CHOL20_ (chain A) and ZIKV_CHOL40_ (Chain B) systems (Figure S11c and S12c). The aromatic residues Y122 and F123 are in proximity to membrane-water interface and hence remain solvated in all ZIKV systems.

Similar to ZIKV systems, the insertion profiles of DENV NS1 loops were calculated and are shown in Figure S11b and S12b. The residues 118-125 of DENV_POPE_ chain A remains buried beneath the phosphate headgroup, while its chain B shows less insertion with the exception of A117 and W118. Similarly, the residues 118-123 of DENV_CHOL20_ chain A stays close to the headgroup, while the chain B remains on the membrane surface (Figure S11b). The intertwined loops which stay close to the head group and inserted into the membrane are highly probable to adopt a secondary structure (Figure S10) The DENV intertwined loop hydration profile was calculated and is shown in the Figure S11d and S12d (shown in dark green, light green and orange). It is observed from the secondary structure and insertion profile analysis that the residues 118-125 are forming a secondary structure and stay close to head group than rest of the loop residues. Similar to ZIKV systems, the polar and charged (basic) residues such as K116, K120, S125 are more hydrated, with the exception of T117. The residue G119 attracts few waters in all 3 systems, perhaps due to lack of side chain with functional groups that can interact with water. The residue L124, which is inserted in DENV_POPE_, shows a non-uniform water distribution (Figure S11d and S12d). For example, in the chain A of DENV_POPE_, it lies below the headgroup and therefore attracts less water, whereas in the cholesterol containing complexes (DENV_CHOL20_, DENV_CHOL40_) it attracts more waters due to solvent exposure. The variant aromatic residues Y122 and F123 of ZIKV are substituted respectively by the basic and hydrophobic residues K and M in DENV NS1 at the structurally equivalent positions. Consequently, the residue K122 attracts more waters in chain B of all three DENV complexes (Figure S11d and S12d).

Altogether, the secondary structure of the disordered intertwined loop residues 118-126 is influenced by their membrane binding property. A possible secondary structure can be seen when the loop interacts favourably with the membrane and expresses stable interactions. According to the insertion of individual residues, their hydration state is also affected. Consistent with the density plots, ZIKV NS1 shows more insertion than DENV NS1 possibly due to the substitutions like S121A, Y122K, F123M and R125S in DENV. Altogether, our study shows that the distinct domains of NS1 binds distinctly and insert well into the lipid bilayer. It is also clear that the NS1 shows distinct binding dynamics with different cholesterol concentrations in the bilayers. To unveil how the different lipid bilayers modulate the NS1 binding, their interaction energy and contact analysis were analyzed further.

### Role of Electrostatics in regulating NS1 interaction

The van der Waals and electrostatic components of interaction energies between the NS1 protein and POPE/Cholesterol were calculated for the last 300 ns of simulation and the average interaction energies are reported in Table S2. The total interaction energies between NS1 and membrane were calculated by summing up the van der Waals and electrostatic energy terms with a cut-off distance of 12 Å. The average interaction energy of ZIKV systems is more favorable than that of DENV systems, and the presence of cholesterol increases the stability of NS1 binding. In particular, electrostatic energy dominates the binding between NS1 and POPE, while van der Waals dominates the binding between NS1 and cholesterol in both the ZIKV and DENV systems. The electrostatic surface potential will help to understand this difference in interaction energy. Hence, the electrostatic potential was generated for the structures extracted at the end of 600 ns simulation.

The electrostatic potential was calculated using the APBS package^37, 38^ and the generated potential plotted on the NS1 isosurface is shown in Figure 6. The inner face of the NS1 was analyzed as it binds to the membrane. The inner face of NS1 consists primarily of the β-roll, a portion of the wing (connector, intertwined loop), and the C-terminal domains. These regions exhibit sequence diversity between ZIKV and DENV (Figure S2). Figure 6 depicts basic and acidic residues in ball and stick representation of crystal structures (Figures 6a and 6f), with the electrostatic potentials plotted on the surface of the ZIKV and DENV NS1 β-roll and wing domain.

**Figure 6:**
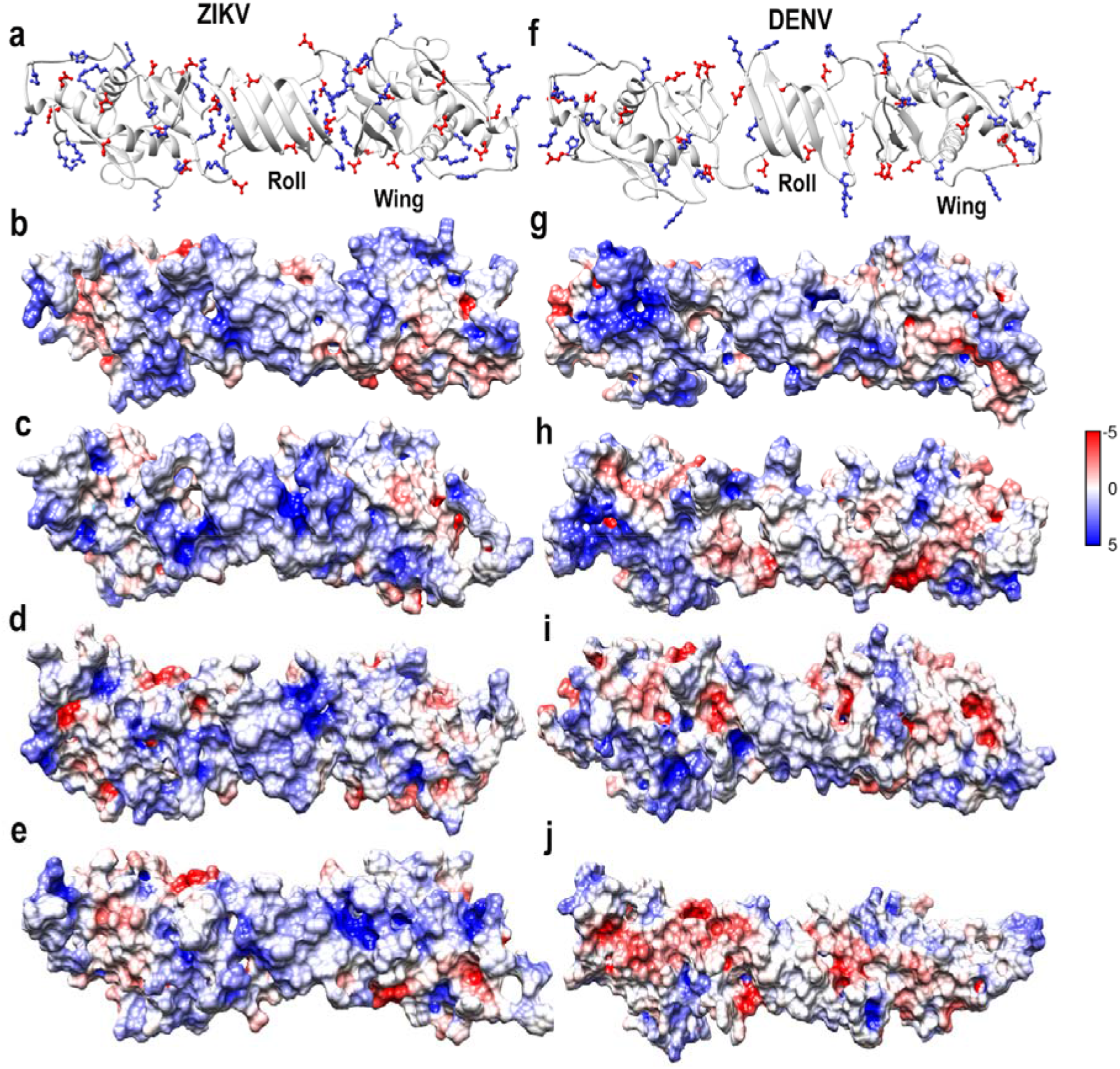
The inner face of (a) ZIKV and (f) DENV NS1 crystal structures consisting of β-roll, wing shown in ribbon representation. Acidic (red) and basic (blue) residues are shown in stick representation. Electrostatic potential maps of ZIKV (b-e) and DENV (g-j) NS1 proteins calculated for the same region for apo (b, g), POPE (c, h), cholesterol 20% (d, i) and cholesterol 40% (e, j) containing bilayers depicting the negative to positive potentials in the color scale of red to blue. The protein is shown as cartoon with basic and acidic residues in ball and stick, colored in blue and red respectively. Colors scale range of –5 to 5 mV.

It is evident from the sequence in Figure S2 that the ZIKV NS1 contains more basic residues than DENV. Accordingly, the ZIKV NS1 predominantly shows blue patches than DENV NS1. The electrostatic potential of apo NS1 shows distinct variation between ZIKV and DENV NS1. In the ZIKV_POPE_, several basic patches are seen, especially in the wing domain (Figure 6c). On the other hand, DENV surface shows both basic and acidic patches (Figure 6h). In the presence of cholesterol, the basic patches are retained in the ZIKV NS1 surface (Figure 6 e,f) but in the DENV structures the acidic patches are enhanced. In particular, DENV_CHOL40_ shows more acidic patches in the wing domain (Figure 6j). Thus, the pattern of electrostatic potential between ZIKV and DENV are very distinct and hence, the residue-level interactions might shed further insights on the sequence-dependent variation in the interaction.

### Interface of the NS1 and membrane interaction

The primary source of NS1 protein’s interactions with the lipid bilayer is through the inner face which was established from the above analysis. Our density plot and interaction energy analyses have provided enough evidence that NS1 exhibits varying degrees of insertion in the simulated systems and that the stability of NS1-membrane interactions is dependent on the electrostatic interaction. Prior research has identified the β-roll and finger loop of the wing domain as the key membrane-interacting regions ^19-21^. Additionally, our study identifies the role of intertwined loop and C-terminal domain in membrane binding. To investigate the interaction of NS1 with various membrane components, the lipid heavy atoms within 4 Å of each protein residue which remain in stable contact for at least 90 ns of the last 300 ns simulation are listed (Tables 2 and 3) for both ZIKV and DENV systems. The number of residues is classified based on the type of amino acids such as acidic, basic, aromatic, hydrophobic residues and lipid groups (headgroup and acyl group). The cumulative number of residues forming contact for both the chains are listed in Tables 2 and 3.

**Table 2.** Residues of ZIKV NS1 protein showing stable contacts with different lipid components^1^.

| Residues | ZIKV<sub>POPE</sub> | ZIKV<sub>CHOL20</sub> | ZIKV<sub>CHOL40</sub> |
| --- | --- | --- | --- |
| **Lipid Head group** |  |  |  |
| Acidic | Chain A: D30, D157, D12, E26, E74, E80, E203, E274<br>Chain B: D30, E26, E74, E192, E272, E274, E278 | Chain A: D7, D30, D12, E26, E74, E80, E81, E272, E274, E278, E279<br>Chain B: D1, D30, D37, D157, D12, E26, E74, E80, E272 | Chain A: D7, D30, D157, D327, D12, E26, E74, E81, E192, E272, E274, E278, E279, E309<br>Chain B: D30, D37, D157, E26, E74, E272 |
| Basic | Chain A: K11, K33, K116, K120, R14, R29, R31, R214, H164<br>Chain B: K33, K120, R14, R29, R31, R40, H164 | Chain A: K11, K33, K326, R29, R31, R40, R214, R276, R324, H164, H216<br>Chain B: K11, K33, K116, R14, R29, R31, R40, R125, H35, H164 | Chain A: K10, K11, K33, K213, K326, R14, R29, R31, R40, R125, R214, R276, R306, H164<br>Chain B: K11, K33, K120, R14, R29, R31, H158, H164, H216 |
| Aromatic | Chain A: F123, F163, Y122, W28, W115, W118<br>Chain B: F160, F163, W28, W168 | Chain A: F8, F160, F163, W28<br>Chain B: W28, F160, F163, Y122 | Chain A: F123, Y122, W28, Y122<br>Chain B: W28, F160, F163 |
| Hydrophobic | Chain A: T13, C15, G16, T17, A27, A117, G119, V124, G161, V162, S121<br>Chain B: C15, T17, A27, S70, G159, G161, V162, T165, V162 | Chain A: S9, T13, C15, T17, G305, G16, A27, G161, V162, P281, A325,<br>Chain B: T13, C15, G16, T17, A27, G73, A77, S121, V124, G161, V162 | Chain A: S9, T13, C15, S70, S121, A27, P39, A77, I78, G159, P281, G282, T283, G305, V307<br>Chain B: C15, N23, A27, S70, G73, A77, G119, G159, G161 |
| **Lipid Acyl chain** |  |  |  |
| Acidic | --- | 1D | --- |
| Basic | Chain A: K11, R29, R31<br>Chain B: | Chain A: K11, R29<br>Chain B: R29, R125, H164 | Chain A: K11, R14<br>Chain B: R29 |
| Aromatic | Chain A: W118,<br>Chain B: F160, W28, W118 | Chain A: F8, W28,<br>Chain B: F123, F163, Y122, W28 | Chain A: F20, W98,<br>Chain B: F160, Y122, W28 |
| Hydrophobic | Chain A: C15, G16, A117, V162, T13,<br>Chain B: C15, V162 | Chain A:<br>Chain B: V2, C15, V124, V162 | Chain A: S6, G119, S9, T13, C15, S121,<br>Chain B: G119 |

<sup>1</sup>The NS1-lipid contacts are classified according to the amino acid types and the lipid groups. The phosphate and the glycerol backbone of lipids were together considered as head group, while the contacts with acyl chains are tabulated separately. A contact is defined by a distance cutoff of 4 Å that is stable for at least 30% (90 ns) of the last 300 ns of the trajectory.

**Table 3.** Residues of DENV NS1 protein showing stable contacts with different lipid components^1^.

| Residues | DENV<sub>POPE</sub> | DENV<sub>CHOL20</sub> | DENV<sub>CHOL40</sub> |
| --- | --- | --- | --- |
| **Lipid Head group** |  |  |  |
| Acidic | **Chain A:** D92, D278, D281, D327, E30, E127, E274, E326<br>**Chain B:** D1, D157, E12, E30, E74, E81 | **Chain A:** D1, D23, D278, D281, D327, E12, E30, E274<br>**Chain B:** D1, E12, D23, D157, E30 | **Chain A:** E12, E30, E37, D23, E276, E278, D281, D327<br>**Chain B:** D12, D23, E30, E74, E326 |
| Basic | **Chain A:** K9, K11, K33, K116, K122,<br>**Chain B:** K14, K116 | **Chain A:** K9, K11, K14, K33, K272, R299, R322, R324<br>**Chain B:** K11, K14 | **Chain A:** K11, K116,<br>**Chain B:** K122, K272, R322, R324, H26 |
| Aromatic | **Chain A:** F163, W8, W115, W28<br>**Chain B:** F160, Y158, W118 | **Chain A:** F160, F163, F279, W8, W28<br>**Chain B:** F160, Y158 | **Chain A:** F160, F163, W168,<br>**Chain B:** F163, Y32, W28 |
| Hydrophobic | **Chain A:** N10, T29, Q31, T117, S125, T126, G161, V162, T164, G325,<br>**Chain B:** S2, L13, C15, G16, S17, G119, | **Chain A:** S2, N10, L13, C15, G16, S17, T27, T29, Q31, G119, G161, V162, T164, N166, G282, T300, G328, C329,<br>**Chain B:** C15, G16, S17, T22, N24, T27, T29, G159, G161, T164, T165, N166 | **Chain A:** N10, C15, G16, S17, G159, V162, T164, N166,<br>**Chain B:** L13, T22, N24, V25, T27, T29, Q31, V162, T164, G325, G328 |
| **Lipid Acyl chain** |  |  |  |
| Acidic | 0 | **Chain A:** D281, E12 | **Chain A** |
| Basic | **Chain B:** K122 | **Chain A:** K11, K14,<br>**Chain B** | **Chain A:** K14,<br>**Chain B:** H26 |
| Aromatic | **Chain A:** W28<br>**Chain B:** W118 | **Chain A:** F160, W28, W118 | **Chain B:** F163, W28 |
| Hydrophobic | **Chain A:** T29, M123, L124,<br>**Chain B:** T117 | **Chain A:** L13, C15, V162,<br>**Chain B:** S2, L13 | **Chain A:** S2, G3, L13, C15, G16, G161, V162,<br>**Chain B:** T27, T29, Q31, V162 |

<sup>1</sup>The NS1-lipid contacts are classified according to the amino acid types and the lipid groups. The phosphate and glycerol backbone of lipids were together considered as head group, while the contacts with acyl chains are tabulated separately. A contact is defined by a distance cutoff of 4 Å that are stable for at least 30% (90 ns) of the last 300 ns of the trajectory.

In the presence of cholesterol, the number of residues forming stable contacts between acidic residues and the lipid headgroup in the ZIKV complexes increases (15 in ZIKV_POPE_, 20 in ZIKV_CHOL20_, and 20 in ZIKV_CHOL40_). But a limited number of residues form contacts with acyl group (Likely due to the shorter side-chain length of acidic residues) (Table 2). The participating acidic residues of β-roll domain (1D, 7D, E12, E25, E26, D30), wing domain (D37, D157, E74, E80, E81), and C-terminal (E192, E203, E272, E274, E278, E279, E309, D327) are listed in Table 2. However, no such increase in the number of residues in DENV systems is found due to the fact that the insertion of DENV NS1 is not deeper inside the membrane and the same phenomenon is observed in cholesterol containing systems also. The number of acidic residues forming contact with the headgroup region remained between 13 or 14 in all DENV systems (Table 3). In the DENV complex, the acidic residues of β-roll domain (D1, D23, E12, E30), wing domain (D92, D157, E37, E74, E81, E127), and C-terminal (E274, E276, E326, D278, D281, D327) domain are involved in the interactions with lipid head groups.

In addition to acidic residues, the interaction formed by basic residues with distinct lipid groups in ZIKV and DENV was listed (Tables 2 and 3). The number of residues forming interaction in ZIKV NS1 is significantly larger than DENV complex. In the ZIKV_POPE_, ZIKV_CHOL20_, and ZIKV_CHOL40_ complexes, about 16, 21, and 23 residues, respectively, form stable contacts with the headgroup. There were only 7, 10, and 7 basic residues in the DENV_POPE_, DENV_CHOL20_, and DENV_CHOL40_ complexes, respectively. The basic residues in the β-roll (K10, K11, R14, and R29), wing (K33, R31, H35, R40, K116, K120, R125, H164), and C-terminal (K326, R214, H216, R276, R306, and R324) domains of ZIKV NS1 contribute to the interactions with the lipid headgroup (Table 2). A slight increase in the number of residues forming contacts in the ZIKV_CHOL20_, and ZIKV_CHOL40_ complexes are observed compared to those found for ZIKV_POPE_. Also, the longer side chains of basic amino acids facilitates interaction with acyl groups of the bilayer. Similar to interaction with head group, the basic residues of ZIKV forms higher contacts with acyl groups than DENV basic residues (Tables 2 and 3). The main interacting basic residues of DENV’s are from β-roll (K9, K11, K14, H26), wing (H77, K116, K120, K122), and C-terminal (K272, R299, R322, R324) domains. However, a steady rise in the number of interactions in the presence of cholesterol was not seen, as the number of contacts formed by DENV_CHOL40_ are lesser than that found for DENV_CHOL20_ (Table 3). Our density analysis also indicates that DENV_CHOL40_ is less inserted than DENV_POPE_ and DENV_CHOL20_ systems. The higher number of basic residues found in ZIKV (K10, R29, R31, H35, R40, R125, H164, R276) in the lipid-contacting regions compared to the same regions in DENV explains more contacts found in ZIKV with lipid bilayer.

The number of aromatic residues forming contact with lipid groups in different simulated systems is listed in Tables 2 and 3. Notably, the number of aromatic residues in interaction with the headgroup did not differ significantly between the ZIKV and DENV systems. Nevertheless, in the cholesterol systems, the longer aromatic side chains form considerable number of interactions with the acyl chain groups in ZIKV (4, 6 and 5) than in DENV (2, 3 and 2). The inner surface of NS1 has several hydrophobic residues, particularly in the β-roll domain, finger loop, and intertwined loop regions. Consequently, the number of hydrophobic residues involved in the contact is considerably larger than the basic and acidic residues (Tables 2 and 3). These residues interact significantly with the acyl chain groups of the bilayer due to their longer sidechains. Interestingly, in DENV_CHOL20_ NS1 more hydrophobic residues are involved in contacts with the lipid headgroup than that of ZIKV NS1 hydrophobic residues (Table 3). Perhaps, the hydrophobic part of the sidechain of basic residues replaced these hydrophobic contacts. Overall, the contact analysis gives an idea about the interaction of NS1 with the hydrophobic part (acyl group) of the membrane. So, it is expected for NS1 to form interactions with cholesterol groups present in the hydrophobic part of the bilayer. Also, studies have shown that NS1 interacts with lipid rafts enriched with cholesterol ^21, 22^. Thus, we have included cholesterol in our studies to explore the binding of NS1. The affinity of NS1 towards cholesterol was monitored by calculating, the number of cholesterol molecules within 4 Å of NS1. Figure 7 shows the probability distribution of cholesterol around NS1. Not surprisingly, the increased distribution (ranging from 2 to 8) in ZIKV_CHOL40_ indicates that the number of interacting cholesterols is greater in the 40% cholesterol complex as compared to 20% cholesterol complex.

**Figure 7:**
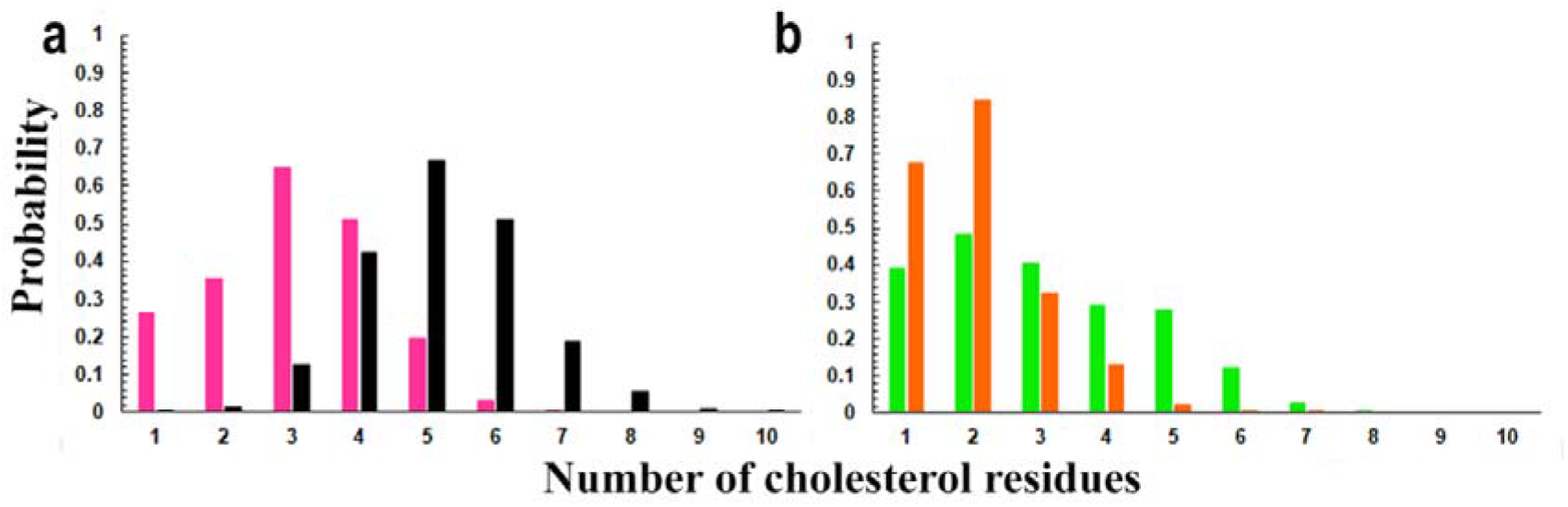
The distribution of cholesterol molecules within 4Å of any residue in ZIKV (a) and DENV (b) NS1 plotted as a probability. The different simulation systems are depicted as ZIKV_CHOL20_ (magenta), ZIKV_CHOL40_ (black), DENV_CHOL20_ (light green), DENV_CHOL40_ (orange).

Around NS1 in ZIKV_CHOL20_, a maximum of 3 to 4 cholesterol molecules are seen, whereas 6 to 7 cholesterol molecules are found to have contacts with ZIKV_CHOL40_ NS1 protein (Figure 7a). The DENV_CHOL20_ complex has 2 to 3 cholesterol molecules (corresponding to the ZIKV_CHOL20_ complex), but the DENV_CHOL40_ system contains just one to two cholesterol molecules (Figure 7b). If we assume the rise in cholesterol around the ZIKV_CHOL40_ is due to an increase in the total amount of cholesterol molecules in the system, we should expect a similar impact for DENV_CHOL40_, which is not the case. Therefore, the rise in the number of cholesterol molecules in ZIKV_CHOL40_ may be the result of higher number of contacts with NS1 and also due to deeper insertion inside the membrane. The analysis reveals both ZIKV and DENV NS1 proteins prefer to interact with cholesterol (at 20% cholesterol concentration) but with increased cholesterol concentration ZIKV alone shows higher interactions.

Altogether, the ZIKV complexes establish more interactions with the bilayer than DENV complexes. Previous experimental studies have reported the role of several residues of β-roll domain (F8, R10 and W28), the conserved intertwined loop residues (W115 and W118) along with other residues (Y122, F123 and V123 in ZIKV; K122, M123 and L124 in DENV) to mediate membrane interactions which corroborates well with our simulations also ^19-21^. Additionally, our study identifies the residues of the intertwined loop of ZIKV (K116, A117, G119, S121, V124, K120, R125) and DENV (K116, K122, A117, G119, L124, S125, T126) to involve in stable contacts with the lipid membrane. A previous mutagenesis experiment on the greasy finger loop residues (residues 159-162) has revealed the important role in membrane binding, which also agrees well with our study ^18^. The membrane interacting residues identified in our study overlap well with the inhibitor-binding site residues reported through molecular docking and virtual screening studies ^29, 30^

## Discussion

Flaviviruses are responsible for human infections that cause illnesses with high rates of morbidity and death. NS1 is a flavivirus non-structural protein that has a variety of roles in viral replication and pathogenesis. Among flaviviral NS1’s many roles, membrane binding is critical for viral replication because it triggers the formation of a replication compartment. The atomic-level interactions and mechanism of membrane binding, however, are yet unknown. The membrane binding properties of the two closely related flaviviral NS1 from ZIKV and DENV are investigated in this work. Using all-atom molecular dynamics simulations, the effect of cholesterol on membrane binding (by varying the cholesterol concentrations 0%, 20% and 40%) and characteristics of the NS1-membrane complex was also investigated. The results of the study reveal that ZIKV NS1 inserts deeper inside the membrane compared to its counterpart in DENV. Electrostatic interactions dominate NS1 membrane interaction, and this is pronounced in the presence of cholesterol. The β-roll, connector regions, intertwined loop, and part of the C-terminal domain were shown to be the most significant membrane interacting regions in our investigation. The contact analysis reveals hydrophobic residues present in the intertwined and finger loop regions. Along with positively charged residues, they drive the membrane interactions supporting the experimental study ^41^. After interaction with the membrane, the dimer NS1 associated with lipid forms the hexameric mNS1. Our results show variation in the membrane binding modes of ZIKV and DENV NS1 proteins.

## Conclusion

Though the structural folds of ZIKV and DENV NS1 homologs are same, their corresponding sequences exhibit ∼50% sequence identity. Our findings reveal that ZIKV and DENV NS1 proteins have different binding preferences for different membrane compositions. Differences in ZIKV and DENV NS1 sequences give rise to variation in their electrostatic potential, which further influences their binding to the membrane. This could be due to the differences in the ZIKV and DENV NS1 sequences. As seen from the results, electrostatic interactions along with the hydrophobic interactions promote the NS1 binding with membrane. The stronger membrane-binding property of ZIKV NS1 could be due to higher basic nature of its inner face than its counterpart in DENV. Contribution of electrostatic interactions for the NS1 binding to the membrane is more pronounced in the ZIKV than DENV. While interacting with the membrane, the structurally and functionally uncharacterized intertwined loop of NS1 was seen to exhibit conformational variability. Furthermore, our results indicate that ZIKV NS1’s intertwined loop insertion into the membrane is deeper more than that found for DENV, and that the presence of cholesterol increases this insertion even more. We have identified key residues necessary for maintaining the NS1-membrane complex from atomic-level interaction analysis, and these findings are consistent with earlier mutational studies. The greater interaction of ZIKV may be responsible for its enhanced remodeling ability of at least 5% than DENV reported in the experimental study^23^. These interacting residues also match the predicted epitope locations on NS1 ^42, 43^, particularly the intertwined loop regions (residues 108-129), the α/β subdomain of wing parts (residues 70-84), the C-terminal domain regions (residues 257-274), and the peptide inhibitor binding sites on the β-roll and C-terminal domain. Despite the fact that the domains of ZIKV and DENV NS1 interacting with the membrane are structurally similar, the nature of their stabilizing contacts differs due to the differences in the residues present in these domains. As a result, the strength of binding with membranes of various compositions also changes. We speculate that variations observed in binding with lipid membranes are likely to allosterically modulate the viral replication process and in turn influence other properties. The findings have implications for the development of possible drugs which will disrupt these interactions (NS1-membrane binding) and design flaviviral specific treatment methods (such as particular antibodies and aptamers).

## Supporting information

Supporting Information with Tables and Figures

## Data and Software Availability

In this study, we have used the experimentally determined crystal structures of NS1 proteins from Zika and Dengue viruses with PDB codes 5K6K and 4O6B. These structures can be downloaded from the Protein Data Bank (https://www.rcsb.org/). Molecular dynamics simulations were carried out using GROMACS ver 5.1 (https://www.gromacs.org/). The minimized coordinates of each system, index files, topology and molecular dynamics .mdp files are available in the GitHub repository (https://github.com/ramasubbu-sankar/zika-virus-NS1-MD). The data (MD trajectories) obtained in this work and the in-house scripts can be obtained from the authors of the manuscript upon reasonable request.

## Acknowledgements

MR would like to acknowledge DBT-RA program in Biotechnology and life sciences for Research Associate fellowship and Institute Post-doctoral fellowship from IIT-Kanpur. RS was Pradeep Sindhu Chair Professor at the time of this investigation. We thank the High-performance computing and Param Sanganak facility at IIT-Kanpur for the computational resources. We thank all our lab members for useful discussion.

## Supporting information

Tables for the structural properties of lipid bilayers, interaction energies between NS1 and lipid bilayer are provided as Table S1 and S2. order parameter of lipid bilayers, pairwise sequence alignment between ZIKV and DENV NS1, RMSD of NS1 without Intertwined loop, conformational landscape of NS1, MD snapshots of ZIKV and DENV, Density profiles of individual domain of NS1 for chain B of ZIKV and DENV, Secondary Structure of intertwined loop residues of ZIKV and DENV, Distance between intertwined loop residues and center of mass of membrane, are presented as Figure S1 to S12. Details of MD simulations of model membrane and bilayers properties are provided.

