## Supporting Information with Tables and Figures for "Membrane-binding properties of NS1 proteins from Zika and Dengue viruses: Comparative simulations in explicit bilayers reveal significant differences"

**Appendix 1**

**Preparation of Model Membranes**

In this study, bilayers of POPE and POPE:CHOL lipid mixtures were built using CHARMM-GUI^1^. Three types of bilayers were used in this study (i) a bilayer with 576 POPE lipids (ii) a bilayer consisting of 550 POPE lipids and 110 cholesterol lipids (20%) and (iii) a bilayer composed of 490 POPE and 196 cholesterol (40%) with lipids uniformly distributed on both sides. The bilayers were solvated with TIP3P^2^ water molecules and treated with CHARMM36 force field^3^. The bilayers were minimized successively using steepest descent and conjugate gradient methods and the systems were equilibrated at 310 K at 1atm pressure. During equilibration, positional restraints were applied on lipid headgroups and were gradually removed in steps of 500 ps. The equilibration was followed by a 200 ns simulation without restraints using the protocol given below. The lipid properties such as area per lipid, thickness and order parameters were monitored throughout the simulation.

**Structural and dynamic properties of lipid bilayer**

The lipid bilayers initially equilibrated for 200 ns and assessed for lipid properties like headgroup area, tail order and thickness of the bilayer. The calculations showed that all the bilayer systems are equilibrated well, and the bilayer properties are monitored in the presence of NS1 and NS1 was manually placed/docked over the equilibrated bilayers.

The lipid headgroup area was calculated for 600 ns time period and shown in table S1. The results showed that the lipid headgroup area of ZIKV_POPE_, ZIKV_CHOL20_, ZIKV_CHOL40_ are (56.0±0.7Å, 52.0±0.5Å and 49.0±0.7Å) respectively. The lipid headgroup area of POPE bilayers in the presence of cholesterol are less when compared to the homogenous POPE bilayers. The thickness of the bilayer was calculated as the distance between the phosphorus atoms of lipids in two leaflets for the last 600 ns simulation trajectories. The calculated thickness of ZIKV_POPE_, ZIKV_CHOL20_, ZIKV_CHOL40_ are 42.08 ± 0.34Å, 44.28± 0.31Å and 45.18 ± 0.29Å respectively. The thickness of the cholesterol containing bilayers increased when compared to the POPE bilayer. The tail order parameters of the two acyl chains of POPE are calculated for the comparing in presence and absence of cholesterol for the last 600 ns simulation time. The tail order parameters for POPE increased in the presence of cholesterol (Figure S1). The bilayers properties of the DENV complexes are calculated and found to be similar to ZIKV complexes.

**Table S1: Structural properties of lipid bilayers**

| **Property** | **POPE** | **POPE+20%**  **cholesterol** | **POPE+40%**  **cholesterol** | **POPE** | | **POPE+20%**  **cholesterol** | | **POPE+40%**  **cholesterol** | |
| --- | --- | --- | --- | --- | --- | --- | --- | --- | --- |
|  | No protein | | | ZIKV | DENV | ZIKV | DENV | ZIKV | DENV |
| Area per lipid (Å) | 56.00±0.06 | 52.00±0.05 | 49.00±0.05 | 56.00±0.07 | 56.00±0.06 | 52.00±0.05 | 52.00±0.04 | 49.00±0.07 | 49.000±0.06 |
| Thickness (Å) | 42.85±0.35 | 44.53±0.36 | 45.65±0.36 | 42.08±0.34 | 42.24±0.39 | 44.28±0.31 | 44.26±0.32 | 45.18±0.29 | 45.26±0.26 |

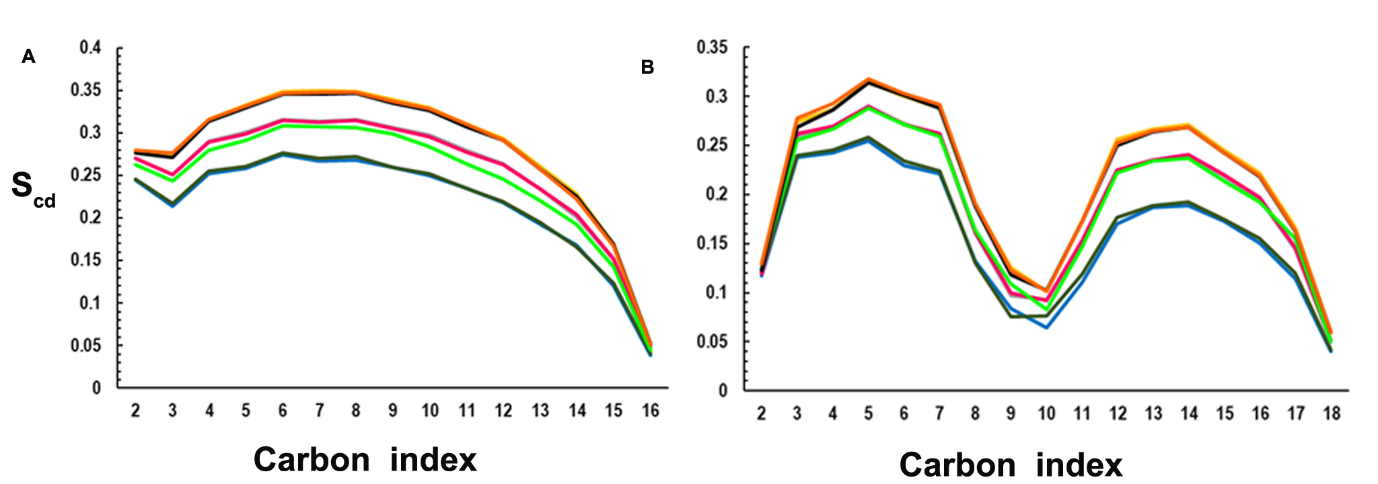

**Figure S1:** Order parameter of the ZIKV_POPE_ (Blue), ZIKV_CHOL20_ (magenta), ZIKV_CHOL40_ (Black), DENV_POPE_ (Dark green), DENV_CHOL20_ (light green) and DENV_CHOL40_ (orange) lipid bilayers. sn-1 of lipid (a) and sn-2 of lipid (b).

**Table S2: Interaction energy between NS1 and POPE and POPE-Cholesterol bilayers, respectively.**

.

| **System** | **Electrostatic Energy (kcal/mol)** | **vdW energy (kcal/mol)** | **Electrostatic Energy (kcal/mol)** | **vdW energy**  **(kcal/mol)** | **Total energy (kcal/mol)** |
| --- | --- | --- | --- | --- | --- |
|  | **POPE** | | **Cholesterol** | | |
| ZIKV_POPE_ | -2677.40±346.1 | -541.98±37.1 | - | - | -3219.39±372.0 |
| ZIKV_CHOL20_ | -3056.88±329.9 | -501.77±42.6 | -18.87±11.3 | -42.19±6.1 | -3613.73±355.3 |
| ZIKV_CHOL40_ | -2944.13±234.0 | -475.85±24.2 | -34.26±14.8 | -96.59±11.6 | -3550.85±238.4 |
| DENV_POPE_ | -2141.52±223.9 | -387.13±30.9 | - | - | -2528.66±233.6 |
| DENV_CHOL20_ | -2498.18±247.1 | -418.76±28.6 | -11.65±10.4 | -34.76±8.1 | -2963.36±260.1 |
| DENV_CHOL40_ | -2206.03±199.8 | -353.62±27.9 | -13.39±13.91 | -33.69±10.8 | -2606.74±207.7 |

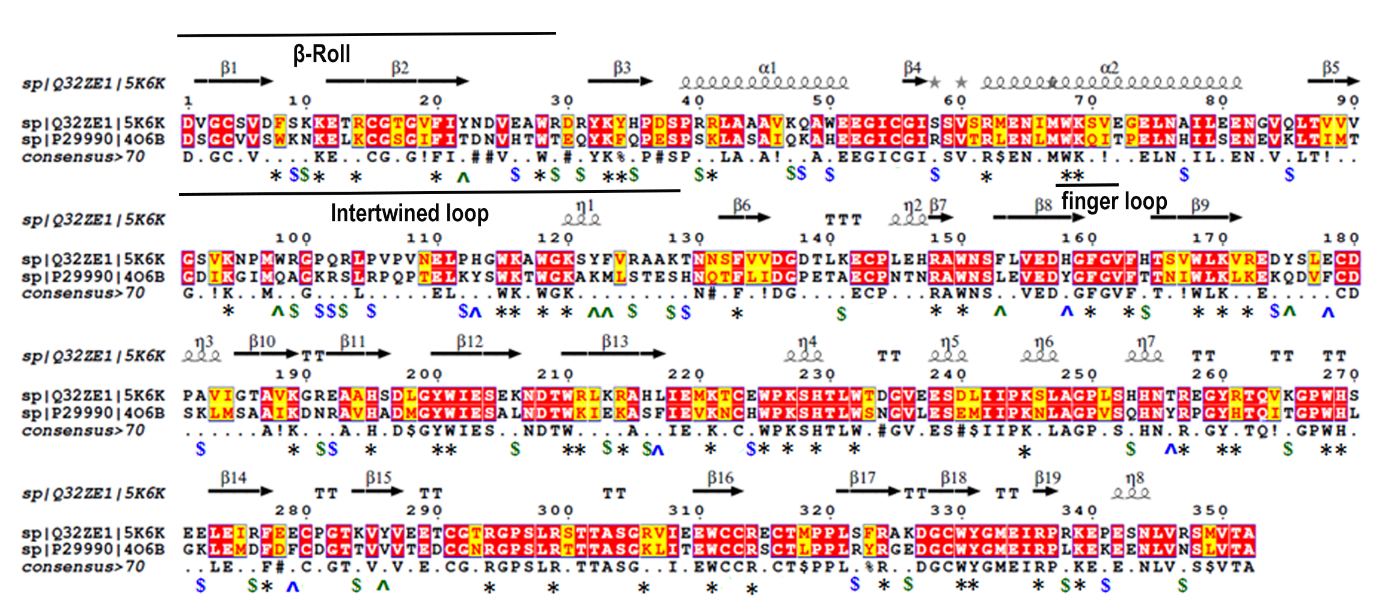

**Figure S2:** Structure-based sequence alignment of ZIKV and DENV NS1 proteins using the ZIKV crystal structure as template representing the conservation of basic and aromatic residues. The variation is marked with ‘$’ and similarity is shown with ‘*’.

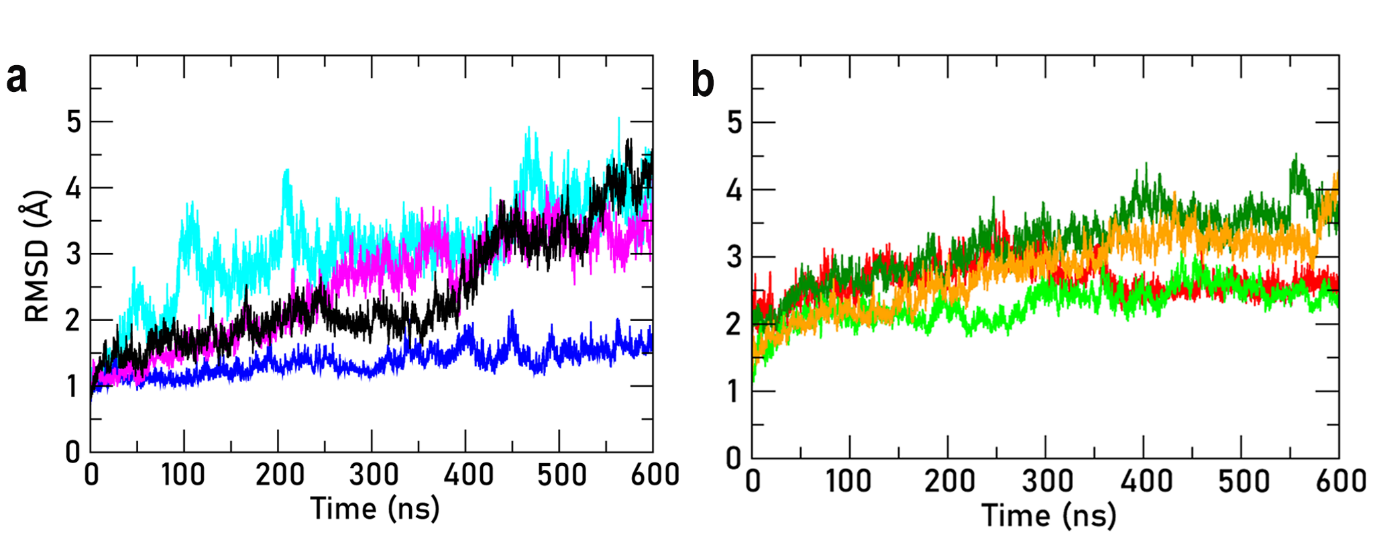

**Figure S3:** Time evolution of the root mean square deviation of ZIKV (a) and DENV (b) without intertwined loop (90-108 a.a). The different NS1 complexes are differentiated as ZIKV_APO_ (cyan), ZIKV_POPE_ (Blue), ZIKV_CHOL20_ (magenta), ZIKV_CHOL40_ (Black), DENV_APO_ (Red), DENV_POPE_ (Dark green), DENV_CHOL20_ (light green) and DENV_CHOL40_ (orange).

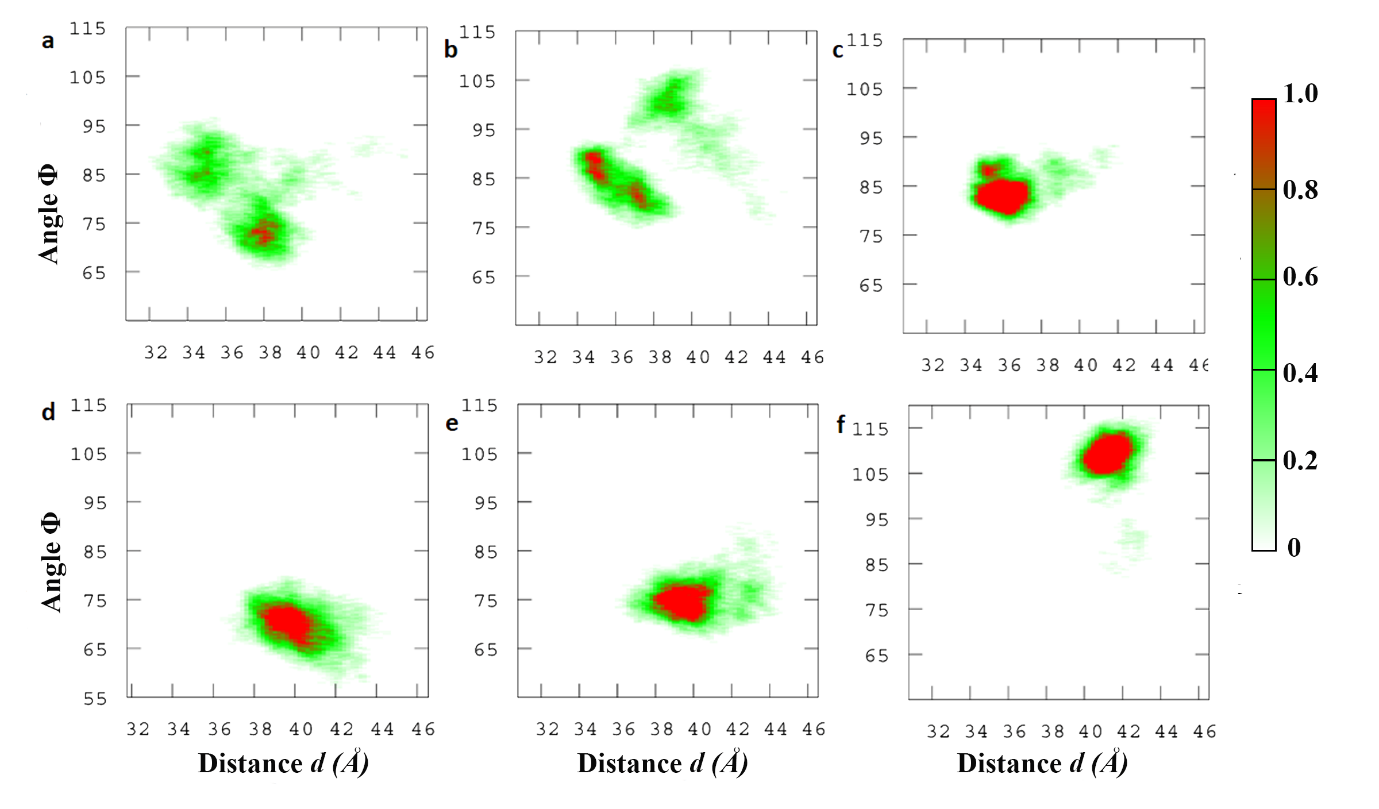

**Figure S4:** The conformational landscape sampled by NS1 as a function of distance d and tilt angle θ for ZIKV_POPE_ (a), ZIKV_CHOL20_ (b), ZIKV_CHOL40_ (c), DENV_POPE_ (d), DENV_CHOL20_ (e), DENV_CHOL40_ (f) are shown

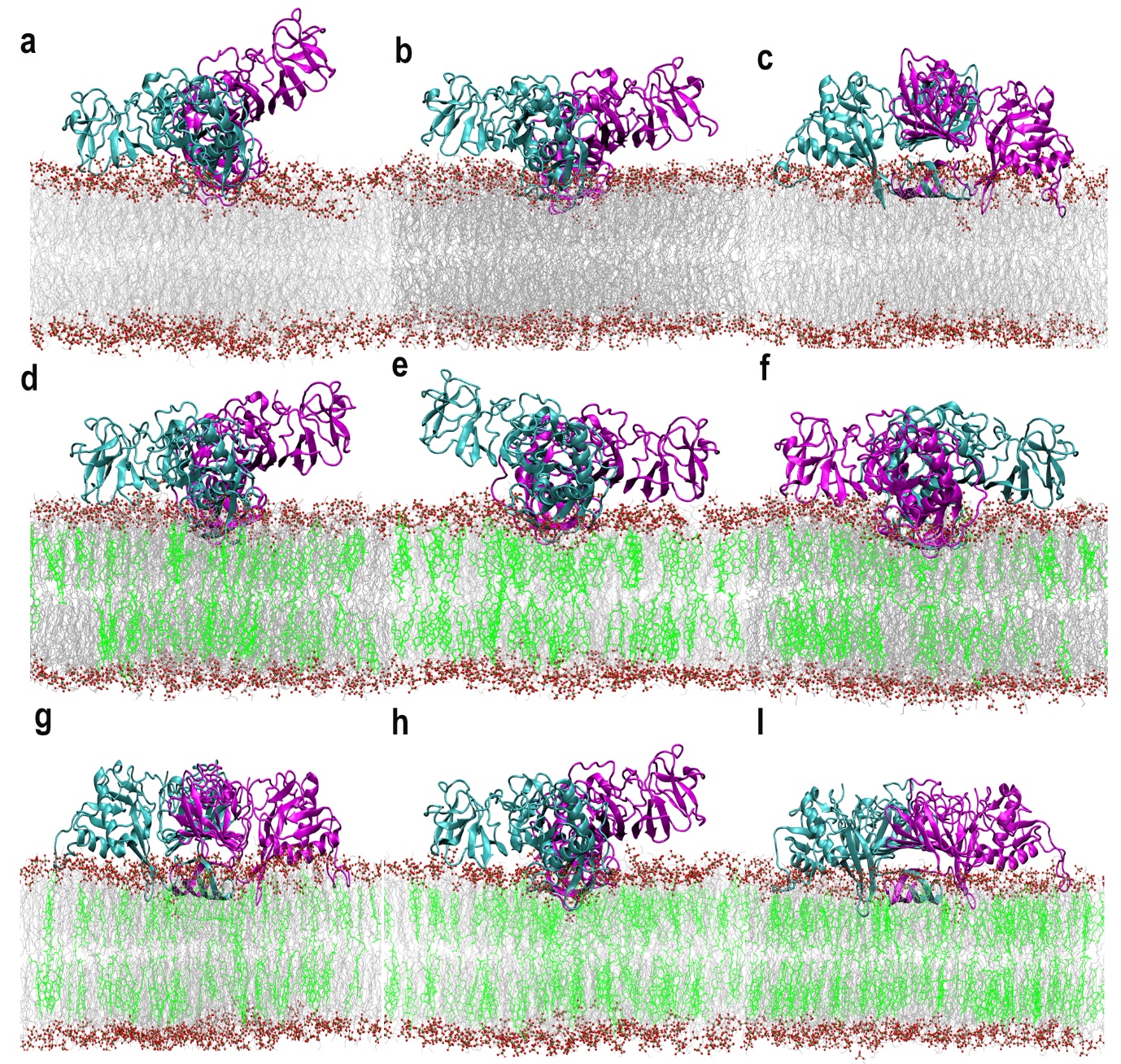

**Figure S5:** Representative structures extracted from each cluster of conformations sampled during the MD simulation of ZIKV_POPE_ (a-b), ZIKV_CHOL20_ (d-g), ZIKV_CHOL40_ (h-i)

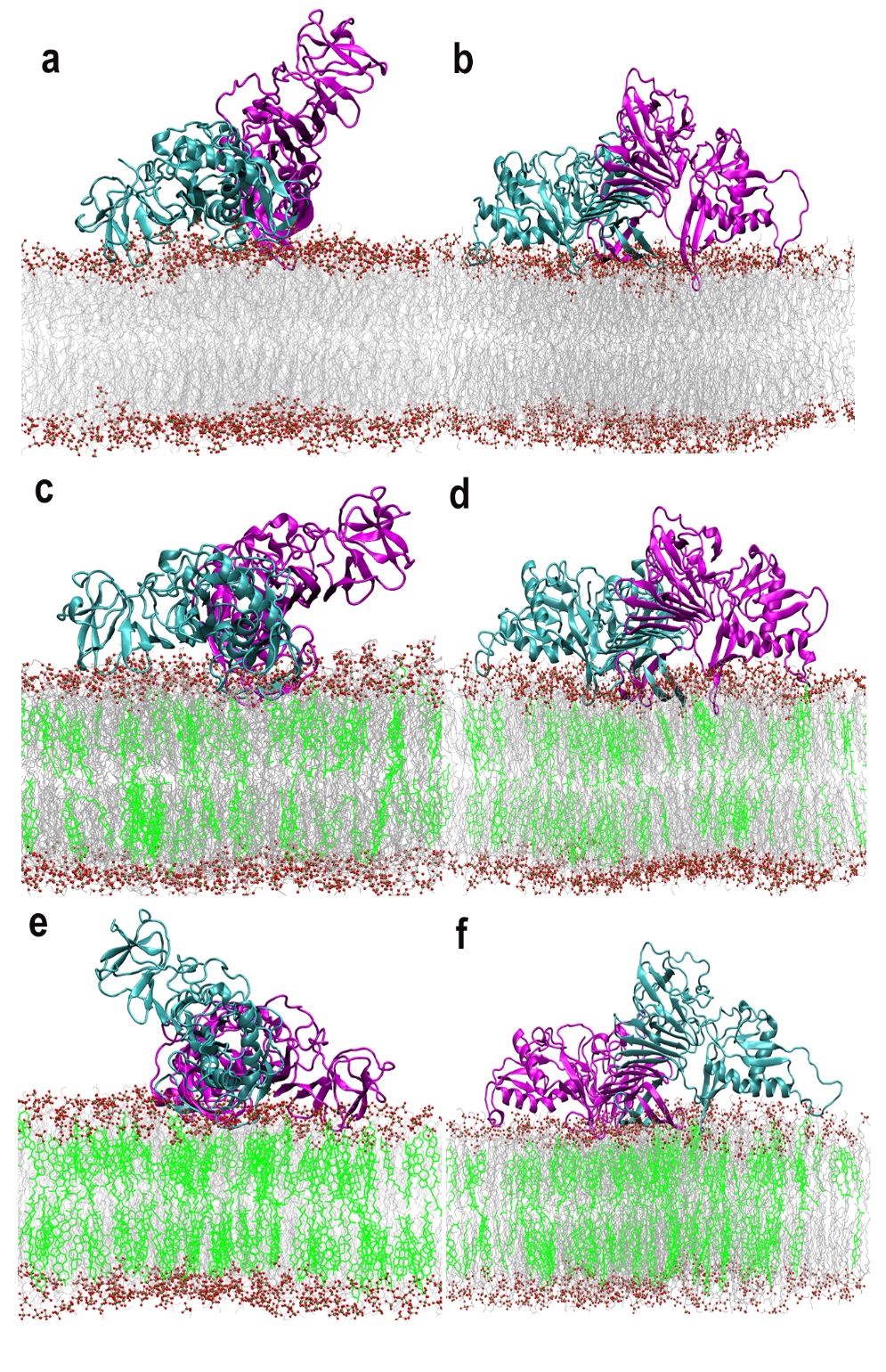

**Figure S6:** Representative structures extracted from each cluster of conformations sampled during the MD simulation of DENV_POPE_ (a,b), DENV_CHOL20_ (c,d), DENV_CHOL40_ (e,f).

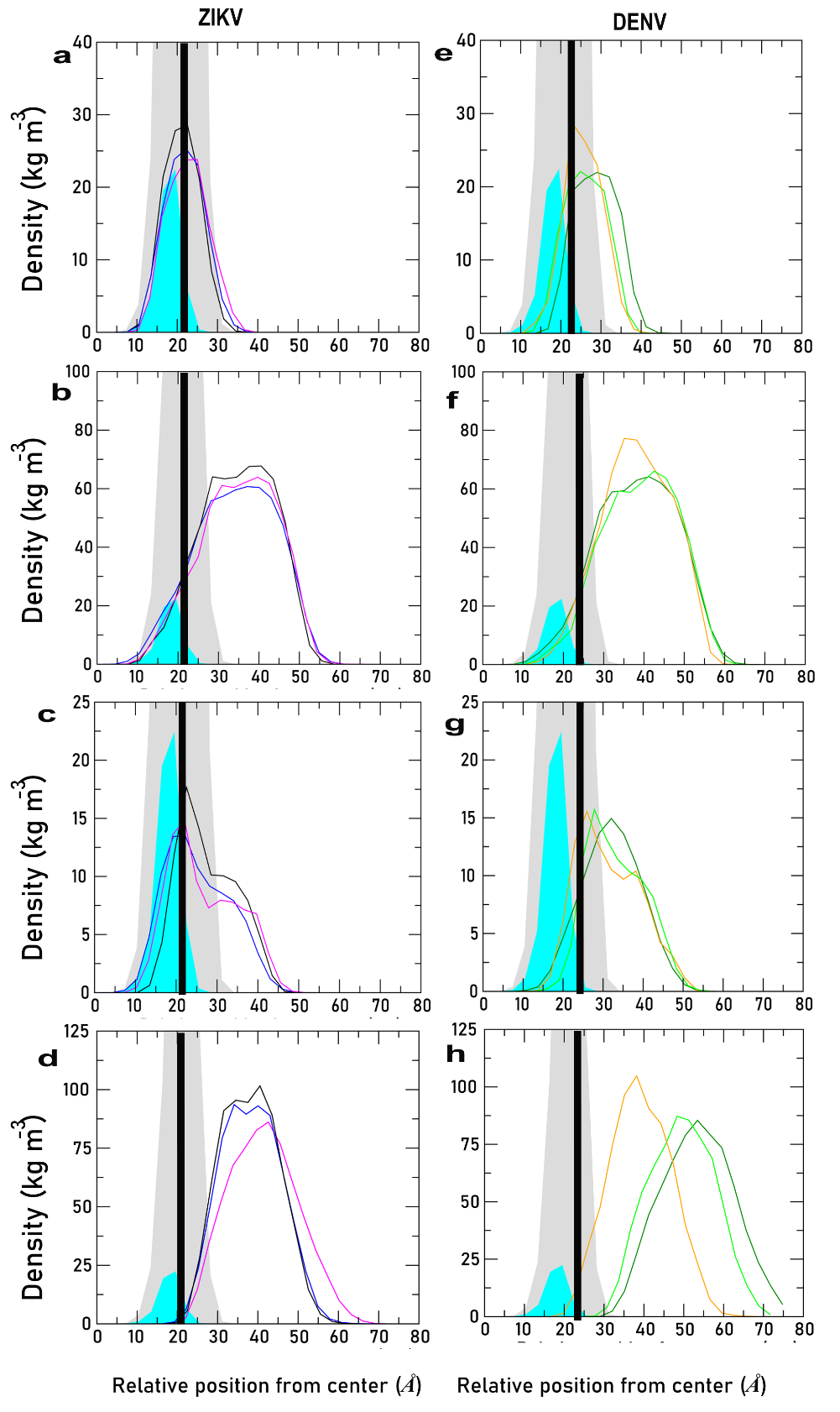

**Figure S7:** Average density profiles calculated for the individual domains of NS1 (Chain B) simulated over the three different bilayers along the Z-axis of the bilayer. The different NS1 complexes are differentiated as ZIKV_POPE_ (Blue), ZIKV_CHOL20_ (magenta), ZIKV_CHOL40_ (Black), DENV_POPE_ (Dark green), DENV_CHOL20_ (light green), and DENV_CHOL40_ (orange). The lipid headgroup region is shown in the grey background with the peak position indicated by the thick black line. The cholesterol density shown in the cyan. The zero on the x-axis corresponds to the bilayer center (as NS1 interacts with top layer and hence only top layer is shown in the figure). The profiles for individual domains are shown in rows labelled β-roll (a,e), Wing (b,f), Intertwined loop (c,g) and the C-terminal domain (d,h) for Chain B of ZIKV, DENV NS1 proteins.

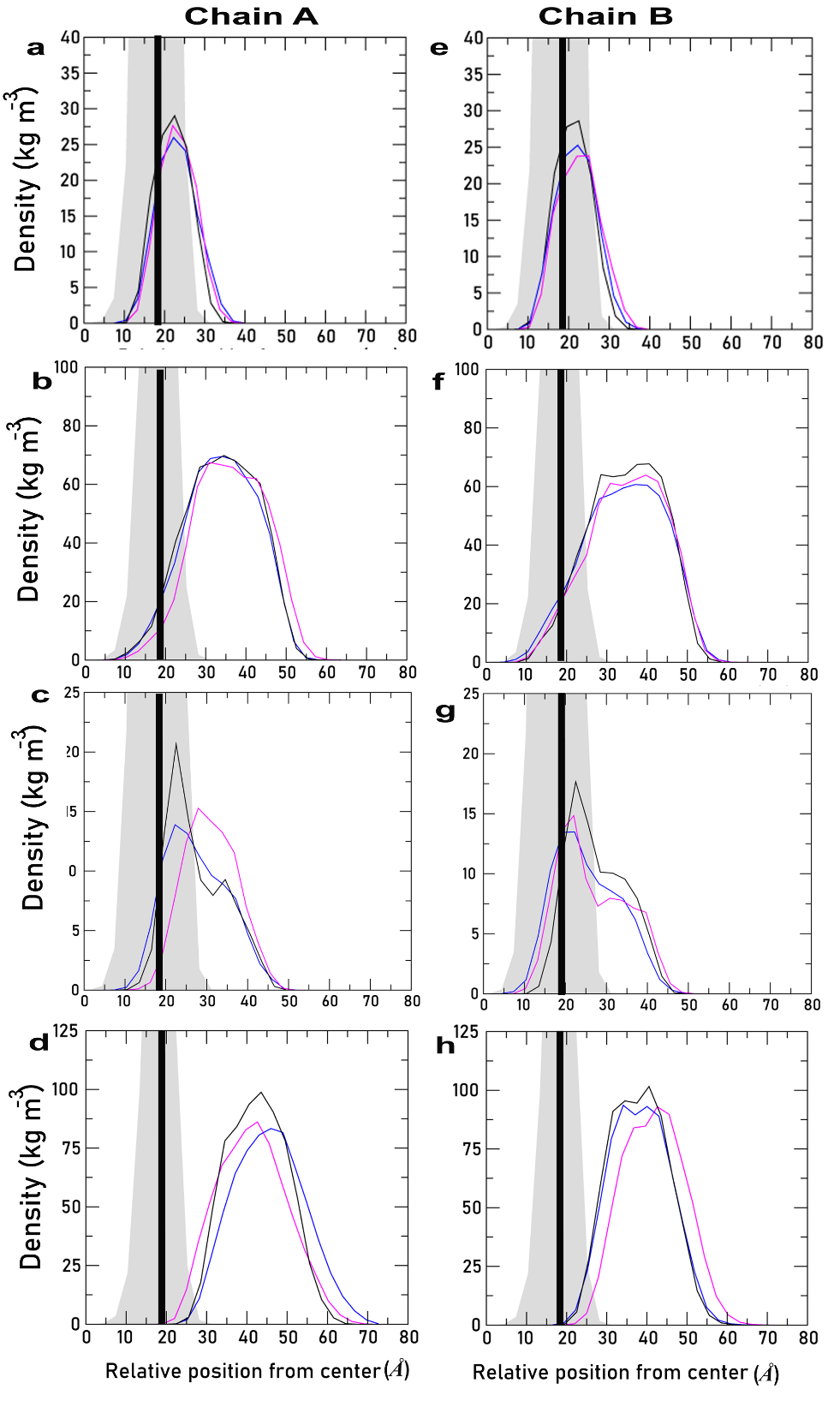

**Figure S8**: Average density profiles calculated for the individual domains of NS1 simulated over the three different bilayers along the Z-axis of the bilayer. The different NS1 complexes are differentiated as ZIKV_POPE_ (Blue), ZIKV_CHOL20_ (magenta), ZIKV_CHOL40_ (Black). The lipid glycerol group is shown in the grey background with the peak position indicated by the thick black line. The zero on the x-axis corresponds to the bilayer center (as NS1 interacts with top layer and hence only top layer is shown in the figure). The profiles for individual domains are shown in rows labelled β-roll (a), Wing (b), Intertwined loop (c) and the C-terminal domain (d). The profiles for individual domains are shown in rows labelled β-roll (a,e), Wing (b,f), Intertwined loop (c,g) and the C-terminal domain (d,h) for chain A and Chain B.

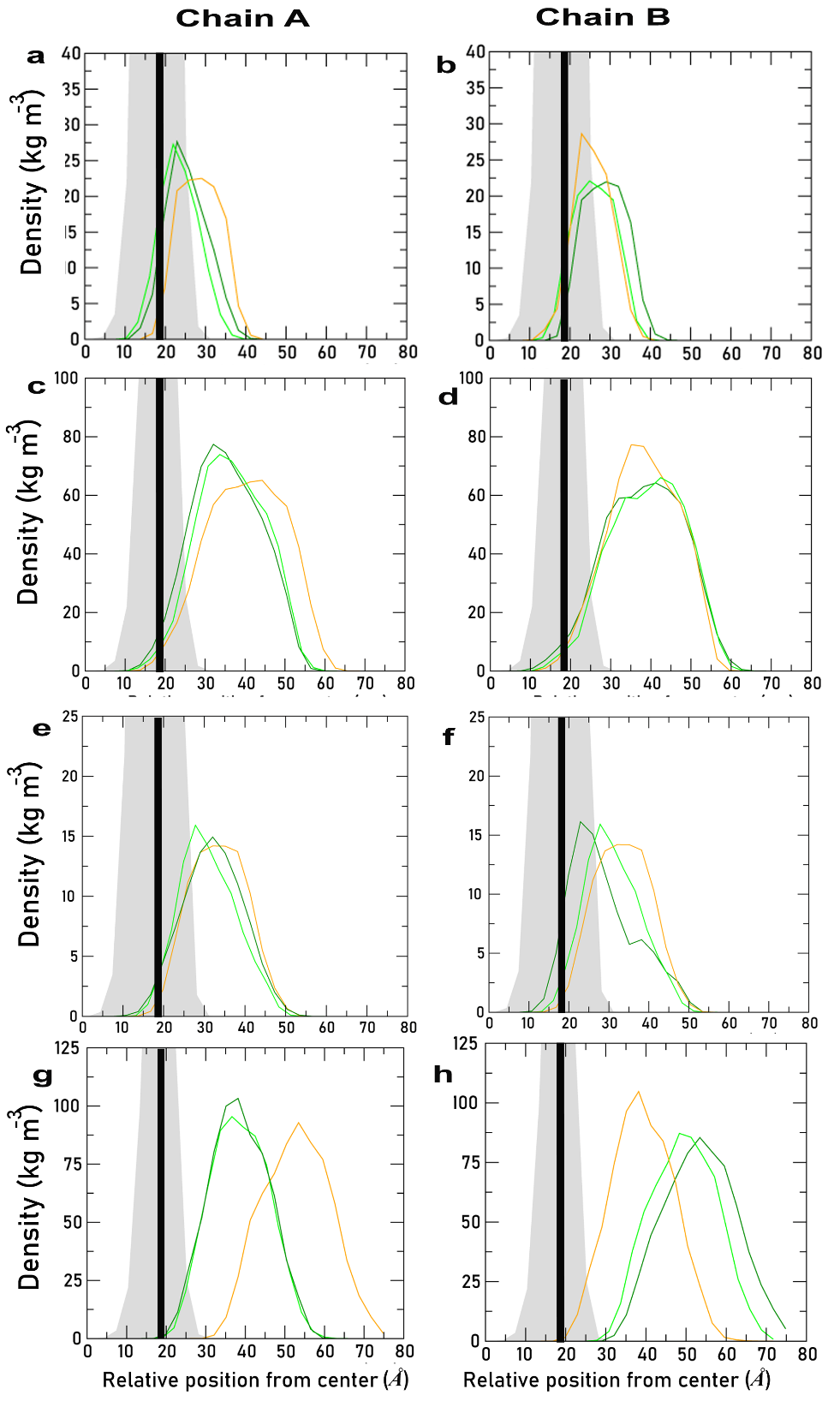

**Figure S9:** Average density profiles calculated for the individual domains of NS1 simulated over the three different bilayers along the Z-axis of the bilayer. The different NS1 complexes are differentiated as DENV_POPE_ (Dark green), DENV_CHOL20_ (light green), DENV_CHOL40_ (orange). The lipid glycerol group is shown in the grey background with the peak position indicated by the thick black line. The zero on the x-axis corresponds to the bilayer center (as NS1 interacts with top layer and hence only top layer is shown in the figure). The profiles for individual domains are shown in rows labelled β-roll (a), Wing (b), Intertwined loop (c) and the C-terminal domain (d). The profiles for individual domains are shown in rows labelled β-roll (a,e), Wing (b,f), Intertwined loop (c,g) and the C-terminal domain (d,h) for chain A and Chain B.

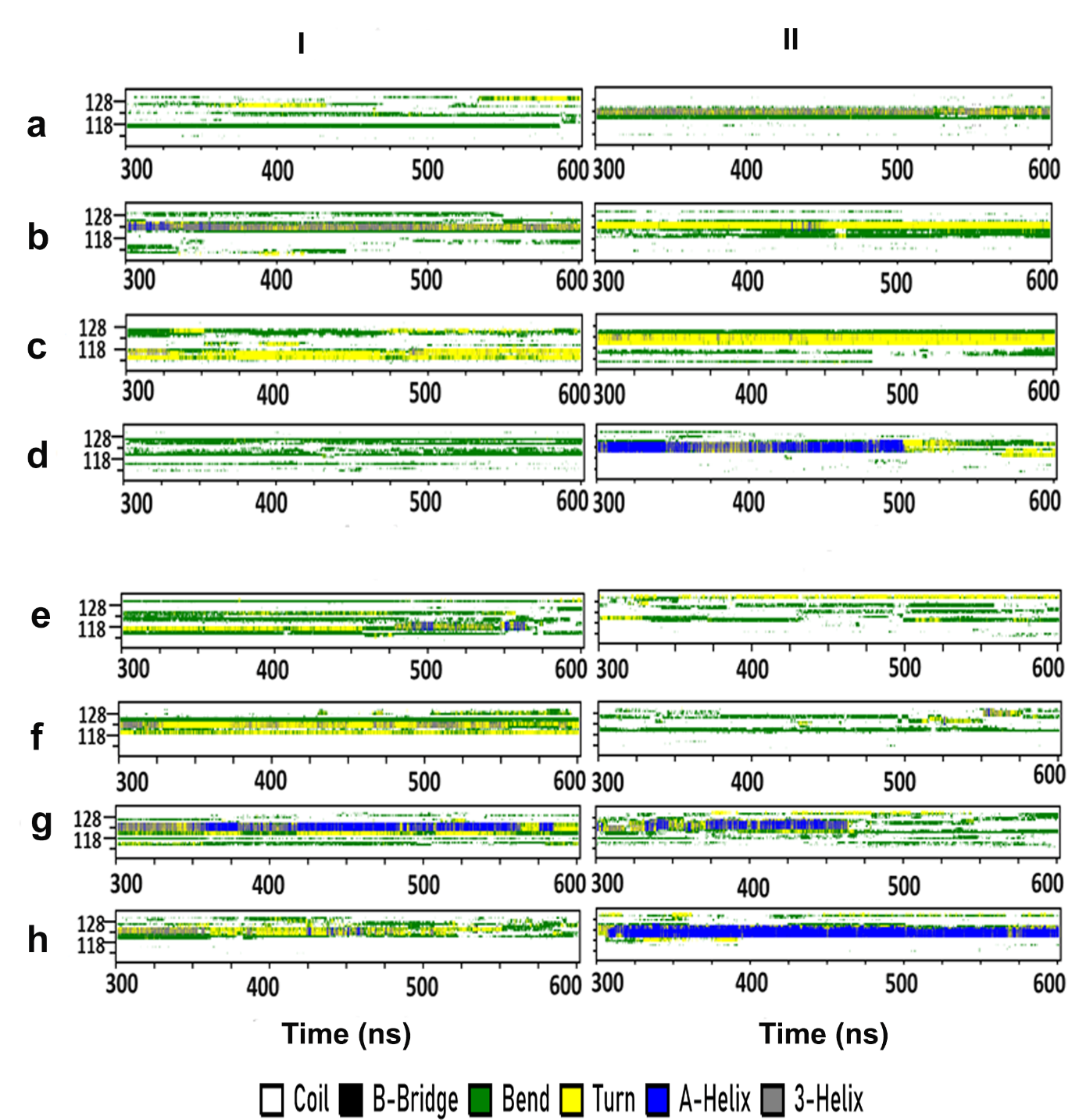

**Figure S10:**Time evolution of the secondary structure elements as a function of time. The panels I (Chain A) and II (Chain B) represent ZIKV_APO_ (a), ZIKV_POPE_ (b), ZIKV_CHOL20_ (c), ZIKV_CHOL40_ (d), DENV_APO_ (e), DENV_POPE_ (f), DENV_CHOL20_ (g) and DENV_CHOL40_ (h).

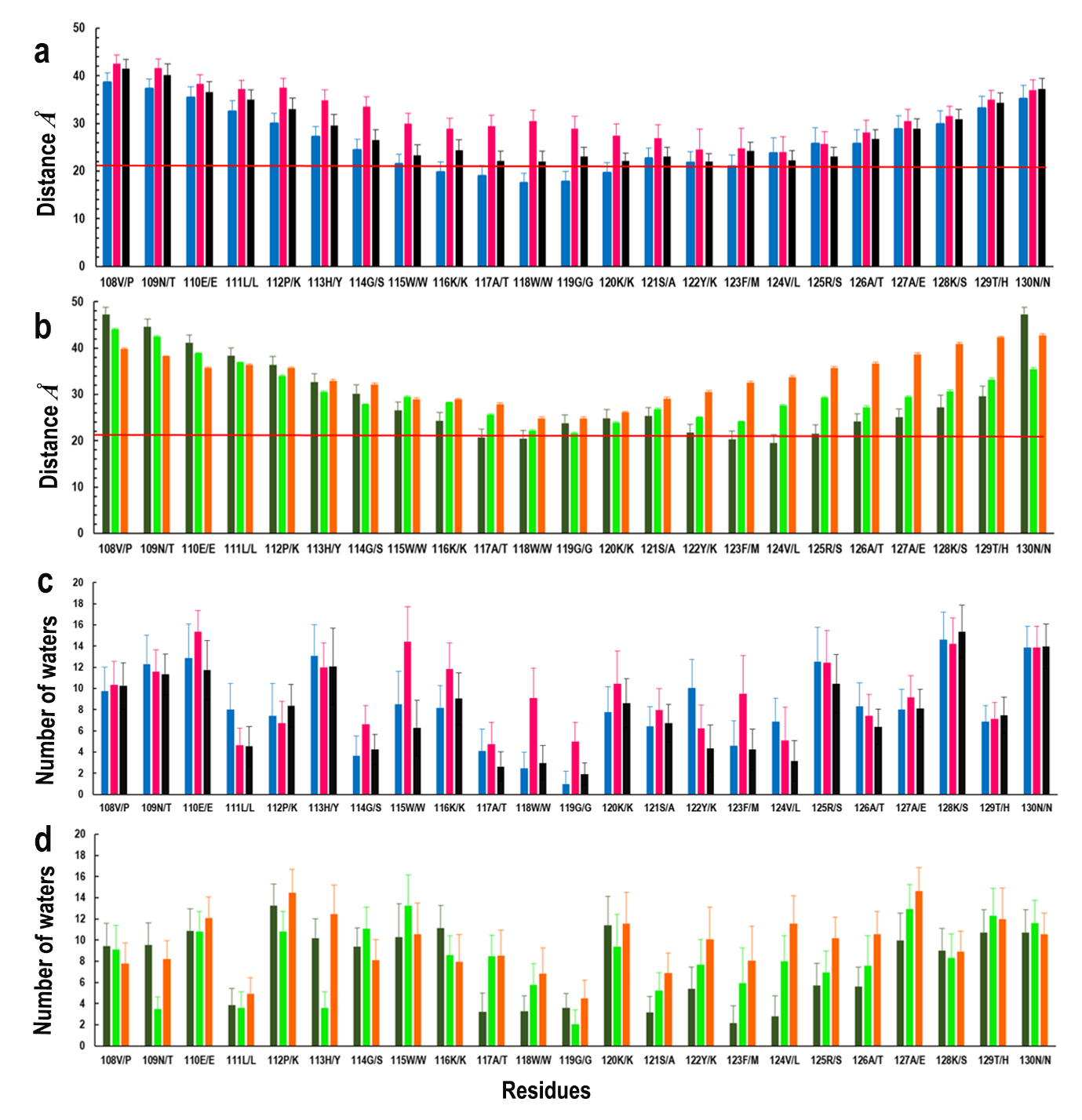

**Figure S11:** The average distance between the center of mass of each residue’s Cα atom (chain A) in the intertwined loop with COM_M_ (a,b) and the average number of water molecules within 4 Å of the residue (c,d). The different simulation systems are colored as, ZIKV_POPE_ (blue), ZIKV_CHOL20_ (magenta), ZIKV_CHOL40_ (black), DENV_POPE_ (dark green), DENV_CHOL20_ (light green) and DENV_CHOL40_ (orange). The horizontal line shown in red color marks the distance between the phosphate head-group and COM_M_.

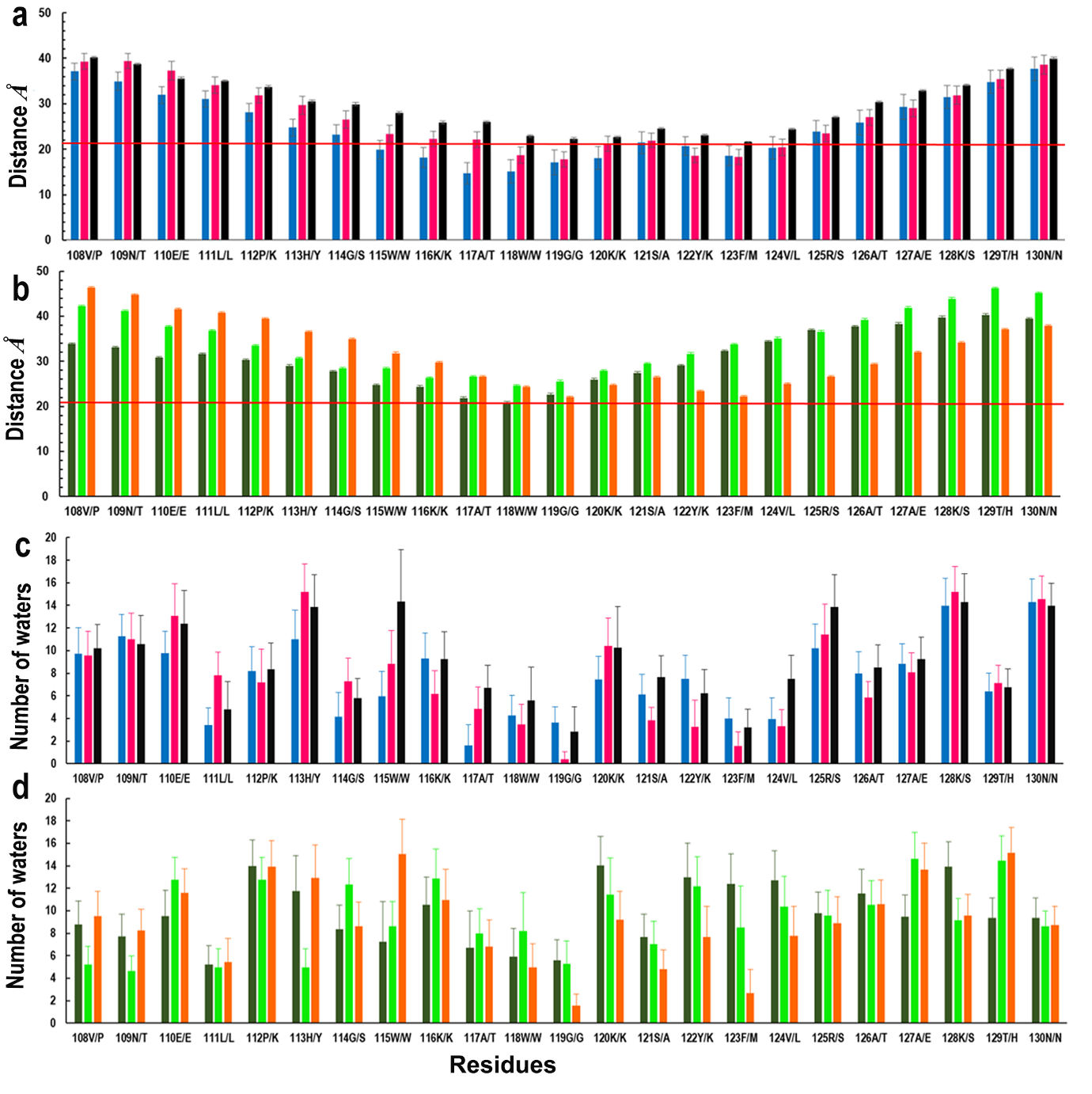

**Figure S12:** The average distance between the center of mass of each residue’s Cα atom (chain B) in the intertwined loop with COM_M_ (a,b) and the average number of water molecules within 4 Å of the residue (c,d). The different simulation systems are colored as, ZIKV_POPE_ (blue), ZIKV_CHOL20_ (magenta), ZIKV_CHOL40_ (black), DENV_POPE_ (dark green), DENV_CHOL20_ (light green) and DENV_CHOL40_ (orange). The horizontal line shown in red color marks the distance between the phosphate head-group and COM_M_.
